# Forward Modelling for Magnetospinography: Systematic Comparison of Boundary Element and Finite Element Methods

**DOI:** 10.64898/2026.06.10.731280

**Authors:** Maike Schmidt, George O’Neill, Meaghan E. Spedden, Stephanie Mellor, Sven Bestmann, Martina F. Callaghan, Gareth R. Barnes

**Affiliations:** Department of Imaging Neuroscience, UCL Queen Square Institute of Neurology, University College London; Department of Neuroscience, Physiology and Pharmacology, University College London, London, United Kingdom; FieldLine Medical, Louisville, CO, United States; Spinal Cord Injury Center, Balgrist University Hospital, University of Zurich, Zurich Switzerland; Translational Neuromodeling Unit (TNU), Institute for Biomedical Engineering, University of Zurich and ETH Zurich, Zurich Switzerland; Department of Clinical and Movement Neurosciences, University College London

## Abstract

Magnetospinography (MSG) enables non-invasive measurement of spinal cord electrophysiology, but accurate interpretation of these signals depends critically on forward modelling assumptions.

We compared four vertebral bone representations: continuous, homogeneous-toroidal, inhomogeneous-toroidal, and MRI-derived-realistic, implemented within both boundary element method (BEM) and finite element method (FEM) frameworks. Lead-fields were computed across the full spinal cord for three orthogonal source orientations. When comparing matched model pairs, BEM and FEM produced consistent forward solutions, with median relative errors below 3.1% and median squared correlation coefficients exceeding 0.998 across all bone geometries. Vertebral bone geometry exerted a systematic and orientation-dependent influence on predicted lead fields. The dominant distinction was between continuous and segmented bone representations rather than between simplified and anatomically detailed segmented models. For left–right oriented sources, segmented geometries produced substantially higher field amplitudes (fT/nAm) than the continuous model throughout the cord (35–72%), while toroidal and MRI-derived realistic models produced comparable results across both frameworks, with median r2 exceeding 0.97 between segmented models compared to 0.67–0.85 for continuous versus segmented comparisons. Sensor placement further modulated sensitivity profiles, with posterior arrays exhibiting greater sensitivity lower down the cord, but anterior and posterior sensors showing comparable sensitivity in the cervical region.

## Introduction

The spinal cord plays a fundamental role in sensory, motor, and autonomic function through ascending and descending pathways that support reflexes, motor coordination, and sensorimotor control^1–4^. Non-invasive access to spinal cord electrophysiology could provide a powerful tool for spinal cord function including neural conduction, pathology, and recovery after injury^5^. However, its white matter tracts are encased by cerebrospinal fluid (CSF) and protected by vertebral bone, forming a complex anatomical and electrical environment^6^. For this reason, along with the small cross-sectional area, deep anatomical location, physiological motion, and surrounding tissue heterogeneity^7, 8^, spinal cord imaging and electrophysiology remain technically challenging^9, 10^.

Magnetospinography (MSG) can be used to address these challenges by measuring the magnetic fields generated by spinal cord activity. Like magnetoencephalography (MEG), MSG benefits from the relative transparency of biological tissues to magnetic fields^11, 12^, enabling non-contact measurements and reduced distortion compared to electrical modalities such as electrospinography (ESG)^13–16^. Sumiya et al^17^ used a purpose-built MSG system with super conducting sensors to measure electrical current flow, on a sub-millisecond time scale, propagating through the cervical spinal cord. This cryogenic system also contains built-in Xray imaging, enabling precise source localisation and sub-millisecond resolution of cervical propagation and spinal cord activity^18–20^, though it does not support concurrent assessment of brain activity.

Optically pumped magnetometers (OPMs) further expand the utility of MSG by providing non-cryogenic, flexible, and anatomically conformal sensor arrays^21–25^. These arrays can be configured to measure from both cord and brain simultaneously^26, 27^and ultimately could be worn during body movement^28^. These advances motivate the development of accurate forward models capable of predicting the magnetic fields generated by spinal cord sources within the electrically heterogeneous torso.

Forward modelling is an essential component of all biomagnetic and bioelectrical source localisation techniques^29, 30^. The forward model defines the mapping from a neural current distribution to the measurable fields or potentials observed at the sensors^31^. In quasi-static electromagnetic regimes^32^, this mapping depends on both primary currents generated by active neurons and secondary currents due to conductivity boundaries. For brain applications, relatively smooth geometry allows simplified, even spherical, approximations^33^. In contrast, the spinal cord’s elongated geometry and high curvature preclude such simplifications, requiring more general numerical techniques such as the Boundary Element Method (BEM) or Finite Element Method (FEM)^31, 34^. While these methods are well established in the MEG and EEG literature^35–38^, their behaviour in spinal cord geometries, characterised by strong conductivity contrasts, complex bone structures, and anisotropic white matter, remains relatively unexplored. Here, we focus on the theoretical foundations and comparative behaviour of BEM and FEM forward modelling frameworks in the context of MSG. We construct a hierarchy of volume conductor models, ranging from simplified homogeneous geometries to anatomically detailed representations, to systematically evaluate where the methods converge and where they diverge.

We additionally introduce `msg-coreg`, an open-source, modular pipeline designed for forward modelling and anatomical coregistration in MSG. Although primarily oriented towards OPM-based spinal magnetometry, `msg-coreg` includes generalised BEM forward-modelling capabilities that can also be applied to cryogenic MSG^17^ and ESG^39^ surface-based electrophysiology. By supporting both personalised and template-based anatomy, it facilitates cross-study compatibility and methodological standardisation.

By clarifying the theoretical behaviour of FEM and BEM forward models and establishing a unified pipeline for BEM modelling across modalities, this work provides a foundation for reproducible source modelling in spinal-cord electrophysiology.

## Theory

### Forward Modelling

Forward modelling in bioelectromagnetism predicts the physical signals (electric potentials or magnetic fields) measured outside the body, generated by neural current sources within the head or spinal cord. Given a model of tissue geometry and electrical conductivity, the forward problem computes a deterministic mapping from neural source activity to sensor measurements.

Neural electrophysiological signals are dominated by low-frequency components (*<* 1 kHz). In this regime, the quasi-static approximation to Maxwell’s equations is valid, allowing electric and magnetic fields to be treated as decoupled in time^29, 31^.

The goal of source modelling is to estimate the neural current distribution **j** from the observed sensor data **y**. The forward model defines this mapping linearly as

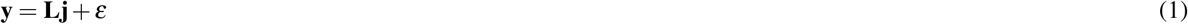

where 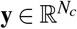 contains the measured magnetic field values (T),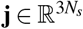 represents the dipole moment amplitudes (Am) at each source location and orientation, 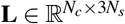 is the lead field matrix, and *ε* accounts for the measurement noise and modelling error. Each column of **L** describes the sensor-level magnetic field pattern produced by a unit dipole at a specific location and orientation. Computing **L** requires solving the electromagnetic forward problem, which we now describe.

For MEG and MSG, each element of **L** is obtained by computing the magnetic field **B**(**r**), at specific sensor locations, from the total current density using the Biot–Savart law:

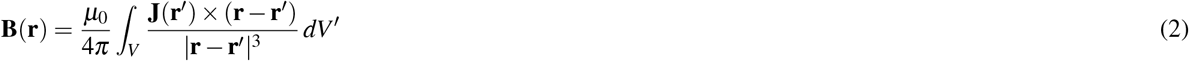

where *µ*_0_ = 4*π* 10^−7^ H m^−1^ is the magnetic permeability of free space. The total current density **J**(**r**) comprises of two components:

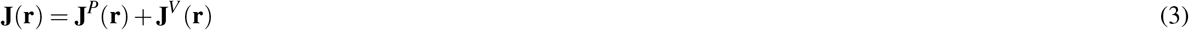

where **J**^*P*^(**r**) (Am^−2^) represents the primary neural currents due to transmembrane ionic flow, and **J**^*V*^ (**r**) is the secondary (volume) current driven by the resulting electric field. Under the quasi-static approximation to Maxwell’s equations, the secondary current takes the form

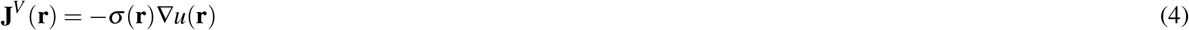

where *σ*(**r**) (S m^−1^) is the electrical conductivity distribution and *u*(**r**) (V) is the electric potential. Charge conservation requires the total current to be divergence-free, ∇· **J**(**r**) = 0, which leads to the governing Poisson equation for the electric potential:

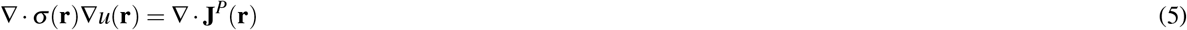

At the outer surface of the conductor *∂* Ω, a homogeneous Neumann boundary condition is applied, enforcing zero normal current flow into the surrounding air: *σ*(**r**)(*∂u/∂n*) = 0 on *∂* Ω where **n** is the outward unit normal.

Neuronal primary currents are modelled as current dipoles

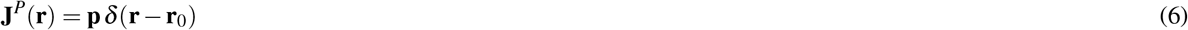

where **p** (Am) is the dipole moment and **r**_0_ is the source location.

Substituting Equations 3, 4 and 6 into the Biot–Savart law (Equation 2), the total magnetic field can be decomposed as

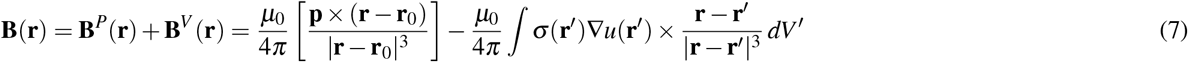

The first term, **B**^*P*^, is the field of a current dipole in an infinite homogeneous medium, an analytically simple expression. The second term **B**^*V*^, captures the contribution of the secondary volume currents, and its evaluation requires knowledge of the electric potential *u*(**r**) throughout the conductor. This is the central computational challenge of the forward problem in MSG, and the primary point of comparison between the numerical methods evaluated in this paper.

Analytical solutions to Equation 7 exist for idealised spherical geometries^34^, but these are poorly suited to the spinal cord due to its elongated shape and complex surrounding anatomy^40^. We therefore employ two numerical methods capable of handling realistic tissue geometries: the Boundary Element Method (BEM) and the Finite Element Method (FEM).

### Boundary Element Method

The BEM reformulates Equation 5 as a surface-integral equation by assuming that the conductor consists of piecewise-homogeneous compartments Ω_*k*_, each with constant conductivity *σ*_*k*_, separated by interfaces *∂* Ω_*k*_. Within each compartment the electric potential satisfies Laplace’s equation, and all conductivity-related effects are confined to the tissue boundaries.

Using Green’s second identity with the free-space Green’s function

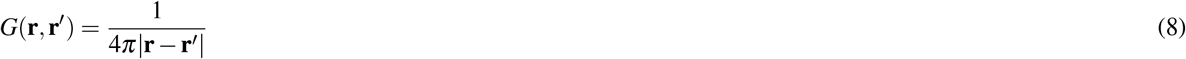

the electric potential at a point **r** can be expressed as a boundary integral:

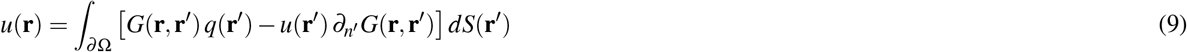

where *q*(**r**′) = *σ*(**r**′) *∂u/∂n*′ is the normal current density (Am^−2^) and *∂*_*n*_′ denotes differentiation along the outward normal **n**′. Continuity of the electric potential and conservation of normal current across conductivity interfaces couple adjacent compartments. Each interface is discretised into surface elements and the boundary fields approximated using piecewise-constant or piecewise-linear basis functions. Enforcing Equation 9 at a set of collocation points yields the linear system

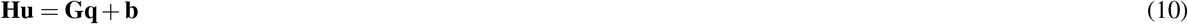

where **u** and **q** are vectors of discretised boundary values, **H** and **G** are dense system matrices arising from the double- and single-layer potential operators, and **b** encodes the primary dipole contribution.

For MEG and MSG, the magnetic field is computed from the solved boundary potentials using the surface-based Biot–Savart law (Equation 2), substituting *q*(**r**′)**n**′ for **J**(**r**′). By restricting computation to tissue boundaries, BEM avoids volumetric meshing and is computationally efficient when tissue compartments can be reasonably approximated as homogeneous.

### Finite Element Method

The FEM solves the governing Equation 5 throughout the full volume without assuming homogeneous compartments. Multiplying by a test function *v*(**r**) and integrating over the domain Ω yields the weak form:

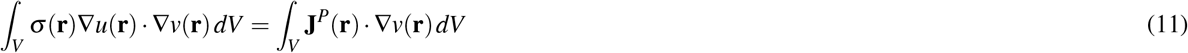

Discretising *u* and *v* using volumetric basis functions defined on a tetrahedral mesh leads to the sparse linear system

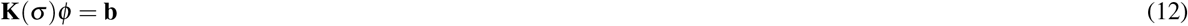

where *φ* contains nodal potential values and **K**(*σ*) is the conductivity-weighted stiffness matrix, a sparse matrix encoding how electrical current is distributed across the mesh given the tissue conductivities assigned to each element. Secondary currents and magnetic fields are then computed from *φ* using numerical quadrature.

FEM naturally supports anisotropic conductivity tensors and complex geometries, making it particularly suitable for spinal cord modelling where cerebral fluid, vertebral bone and white-matter anisotropy strongly influence field propagation. The trade-off is substantially greater computational cost compared to BEM.

## Methods

We evaluated forward solutions for magnetospinography (MSG) using a controlled modelling pipeline based on a single participant-derived anatomical model, four vertebral bone representations, and BEM and FEM solvers. Together, these components enabled systematic assessment of how anatomy, conductivity, and solver choice shape the resulting lead fields.

### Model Construction

A healthy adult participant (male) provided written informed consent under protocol approved by UCL Research Ethics Committee ([3090/005]). A brain-to-whole-spine T1-weighted Magnetic Resonance Image (MRI) was acquired on a 3T Siemens Prismafit scanner equipped with a 32-channel spine coil using a spoiled gradient-recalled echo sequence (1.5mm isotropic resolution, TR = 5.85 ms, TE = 2.10 ms, flip angle = 6°). To enable high quality image data, free of geometric distortion driven by gradient non-linearity, the acquisition was repeated 5 times with the scanner’s bed moved by 15 cm between acquisitions. During scanning the participant lay in a custom-fitted scannercast designed to preserve supine posture between MRI and MSG environments.

Thoracic organs (heart, lungs) were included using meshes from O’Neill et al.^40^ and ECGSim^41^, transformed via a 7-DOF Procrustes alignment followed by a 12-DOF affine ICP refinement. All models were scaled to match a participant-specific optical surface scan (Einscan) captured in the custom-made scannercast, ensuring accurate supine geometry. Outer torso, vertebrae, and spinal cord were segmented from the MRI data with Total Segmentator^42^. Thoracic organs from ECGSim^41^ were transformed into the shared coordinate system and scaled to fit the participant torso (Figure 1).

**Figure 1.**
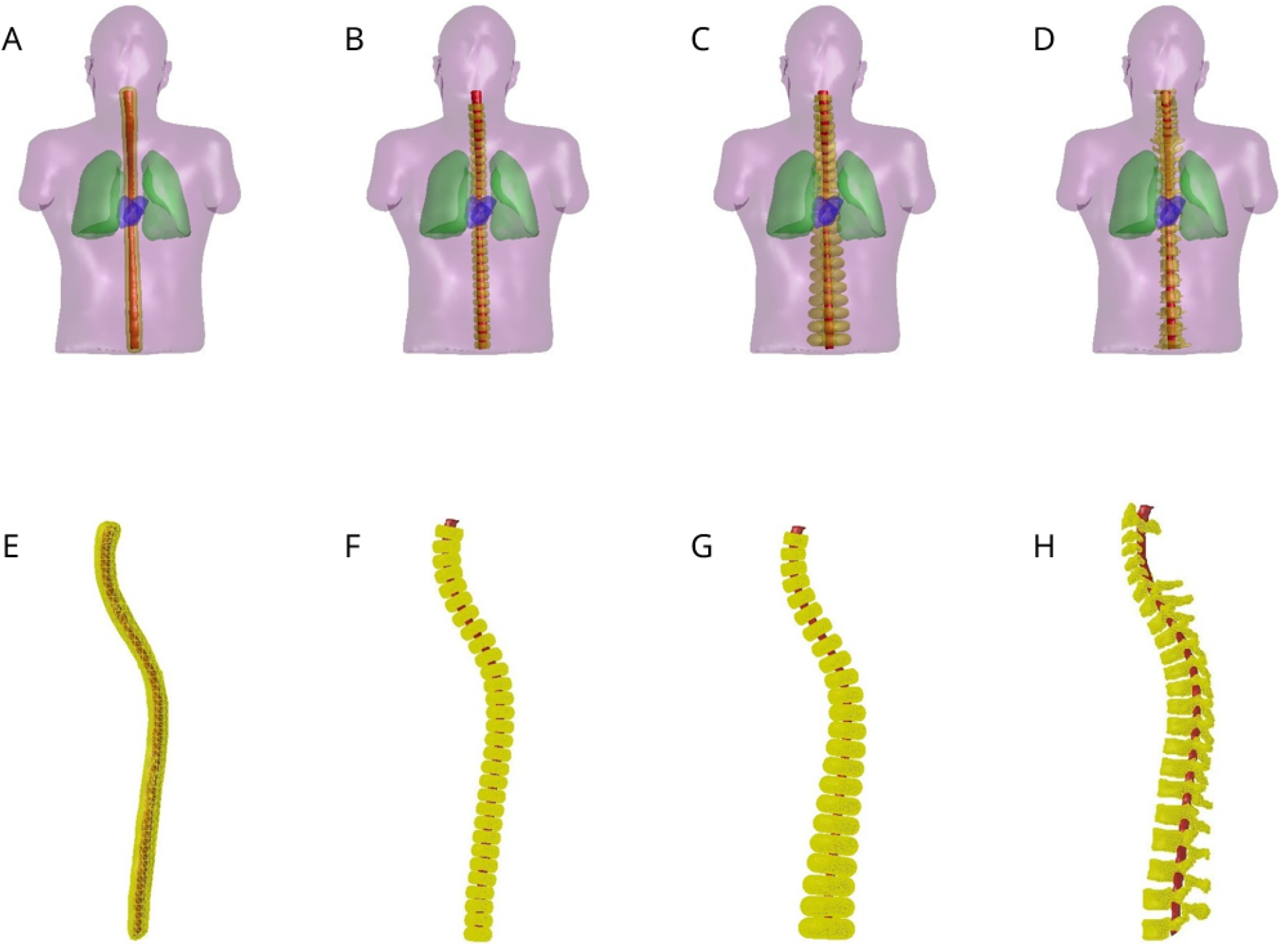
Vertebral bone models and anatomical context. **Top row** (**A**–**D**): each bone model embedded within the participant-specific torso, showing torso (purple), lungs (green), heart (blue), spinal cord (red), and vertebral bone (yellow). **Bottom row** (**E**–**H**): corresponding bone models shown in isolation with the spinal cord (red) included for reference. Models are shown for **(A**,**E)** continuous, **(B**,**F)** homogeneous toroidal, **(C**,**G)** inhomogeneous toroidal, and **(D**,**H)** MRI-derived realistic vertebral geometry.

### Vertebral Bone Model

Four vertebral bone models were implemented (Figure 1, Table 1). All were generated in cervico-thoracic and full-cord versions.

**Table 1.** Vertebral bone model geometry and anatomical compartment mesh characteristics. *Upper section*: vertebral bone models compared in this study. *Lower section*: all anatomical compartments included in forward simulations, with surface mesh resolution and assigned conductivity values. Bone conductivity follows FieldTrip defaults; reported literature values range from 0.004–0.02 S/m^44^

| Compartment | Description | Conductivity (S/m) | Vertices / Faces |
| --- | --- | --- | --- |
| *Vertebral bone models (all assigned $\sigma = 0.00825$ S/m)* |  |  |  |
| Continuous | Normal-based outward extrusion; uniform 12 mm shell | 0.00825 | 3394 / 6784 |
| Homogeneous toroidal | Identical toroidal segments; constant thickness and regular spacing | 0.00825 | 3640 / 7280 |
| Inhomogeneous toroidal | Level-specific scaling; cadaver-derived vertebral dimensions | 0.00825 | 2872 / 5744 |
| MRI-derived | Direct MRI segmentation; full anatomical realism | 0.00825 | 29357 / 58738 |
| *Other anatomical compartments* |  |  |  |
| Spinal cord | MRI segmentation | 0.33 | 5333 / 10662 |
| Torso | MRI segmentation ; 50% mesh reduction applied | 0.23 | 10852 / 21700 |
| Heart | ECGSim mesh, affine-ICP aligned | 0.62 | 1052 / 2096 |
| Lungs | ECGSim mesh, affine-ICP aligned | 0.05 | 916 / 1820 |

The four models respectively capture: (i) a simplified cylindrical shell, (ii) uniformly sized toroidal vertebrae, (iii) anatomically scaled toroids with level-dependent width/height/thickness, and (iv) a fully participant-specific vertebral column.

There are three distinct cross-sectional topologies relative to the spinal cord, with implications for the conductive pathway between a spinal cord source and an external sensor. Figure 2 illustrates representative 2D cross-sections alongside the corresponding conductivity boundaries as represented in the BEM formulation, where surface integrals are evaluated at each interface between a medium with inside conductivity *σ*_i_ and outside conductivity *σ*_o_.

**Figure 2.**
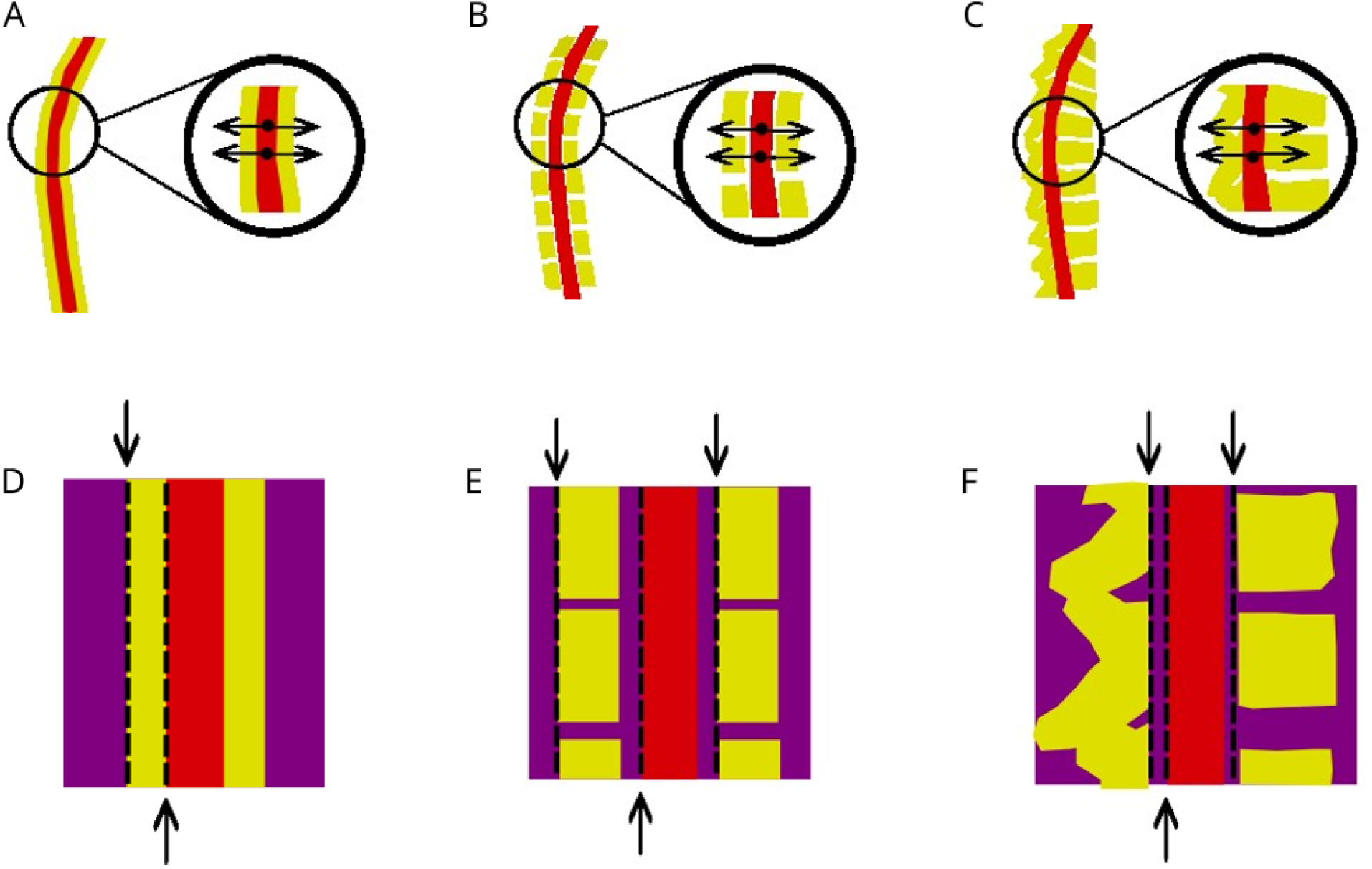
Cross-sectional topology and BEM conductivity boundaries for the three vertebral bone model classes. **Top row (A–C)**: representative 2D cross-sections of the continuous **(A)**, toroidal **(B)**, and MRI-derived **(C)** bone models. The spinal cord is shown in red and vertebral bone in yellow; black arrows indicate representative source locations and their direction towards a sensor array (posterior or anterior). **Bottom row (D–F)**: corresponding conductivity boundaries used in the BEM formulation, where each interface (dashed black line) separates a medium with inside conductivity *σ*_i_ from one with outside conductivity *σ*_o_. The continuous model has two boundaries (cord–bone, bone–torso); toroidal and MRI-derived models introduce a third boundary (cord–torso) at intervertebral gaps. Note that the inhomogeneous toroidal model (not shown here) shares the same cross-sectional topology as the homogeneous toroidal model shown in **(B)** and **(E)**, differing only in level-dependent vertebral scaling.

The continuous model was constructed via normal-based outward extrusion of the spinal cord surface, producing a uniform cylindrical shell with a consistent (cord→ bone→ torso) propagation path and two conductivity boundaries: spinal cord to bone (*σ*_i_ = cord, *σ*_o_ = bone) and bone to torso (*σ*_i_ = bone, *σ*_o_ = torso). The toroidal models were generated by aligning individual toroidal segments to the local spinal cord surface normals. For these models, the conductive pathway depends on rostro-caudal source position: sources within a vertebral segment propagate through bone, whereas sources in an intervertebral gap propagate directly from cord to torso. A third boundary is therefore introduced: the interface between the spinal cord and surrounding torso (*σ*_i_ = cord, *σ*_o_ = torso), with two further surfaces representing the inner and outer faces of the discrete vertebral bone segments (*σ*_i_ = bone, *σ*_o_ = torso). The MRI-derived model was obtained directly from participant-specific segmentation and shares this three-boundary structure, but here the conductive pathway also depends on source orientation due to the asymmetric vertebral anatomy.

To ensure numerical stability and maintain comparability across bone representations and forward modelling frameworks, a uniform surface mesh reduction factor of 50% was applied to the torso model prior to forward modelling. This reduction was consistent with established BEM guidelines recommending coarser discretisation of external boundaries relative to internal tissue interfaces^37^. Systematic investigations of mesh density effects have primarily focused on cortical EEG and MEG models^35, 43^, with limited exploration in spinal cord geometries; we therefore adopted conservative meshing principles consistent with prior MSG work^40^.

Table 1 summarises the mesh characteristics and conductivity values (*σ*_i_) for all anatomical compartments used in the forward simulations. Conductivity values follow the Fieldtrip standard defaults. We note that reported values for bone vary substantially across the literature, with published estimates ranging from approximately 0.004 to 0.02 S/m^44^.

### Sensor Placement and Source Modelling

Anterior and posterior sensor grids were generated by projecting a 30mm rectangular lattice onto the outer torso surface. Rays failing to intersect the torso were discarded. Sensors overlying the face and posterior cranium were excluded to focus on spinal coverage. Magnetically sensitive devices (potentially optically pumped or cryogenic) were modelled as single points elevated by 10mm perpendicular to the local surface, to approximate typical sensor offsets^45^.

Dipolar sources were placed at 5mm intervals along the spinal cord centreline (112 in total), each with three orthogonal 1 nAm dipole moments orientated: left–right (LR), ventral–dorsal (VD), and rostral–caudal (RC). This dipole amplitude is consistent with reconstructed cervical spinal cord sources reported in SQUID-MSG studies, where source amplitudes of 0–2 nAm have been observed following peripheral nerve stimulation^19^.

### Forward Modelling Frameworks

#### Boundary Element Method

Forward fields were computed using the Helsinki BEM Framework (HBF)^33, 35, 43^with linear collocation. Nested surfaces (torso, lungs, heart, spinal cord, bone) defined piecewise-homogeneous compartments. The HBF implementation is openly available at https://github.com/MattiStenroos/hbf_lc_p.

#### Finite Element Method

FEM models were implemented with DUNEuro^46^ using a continuous Galerkin formulation and St. Venant dipole approximation. Volumetric tetrahedral meshes for FEM were generated from BEM surface meshes using Iso2Mesh v1.9 with maximum element volume of 10mm^3^ and minimum dihedral angle of 15°. Resulting meshes, made up of the torso, heart, lungs, spinal cord and bone, contained 106,444 to 144,961 nodes. Tissue-specific conductivities were assigned to tetrahedral elements, and dipole moments were distributed over neighbouring nodes to avoid singularities. We utilised the MATLAB– DUNEuro interface originally written by Medani et al^47^ and modified for this project by O’Neill^40^, available at https://github.com/georgeoneill/study-spinevol/tree/main.

### Comparative Metrics

Lead-field differences between models (indexed 1 and 2) were quantified using relative error (RE) and squared correlation coefficient (*r*^2^)^40^:

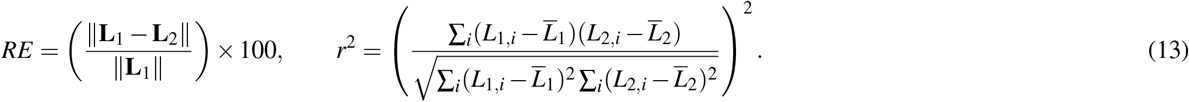

RE captures magnitude and topography differences; *r*^2^ reflects spatial pattern similarity of the magnetic field observed by the sensor array. To provide a consistent reference for pairwise comparisons, the MRI-derived realistic bone model was used as the reference in all cross-geometry comparisons, using the most anatomically realistic as the reference on the grounds that it is a best approximation. Whilst no ground truth forward solution exists, this convention ensures that differences are interpreted as deviations from the most realistic available.

## Results

### Bone Model Comparison

Four vertebral bone representations were embedded within the anatomical torso model: continuous, homogeneous toroidal, inhomogeneous toroidal, and MRI-derived realistic bone. Lead fields were computed across the full spinal cord source space for all dipole orientations. For illustrative purposes, results are presented for a representative thoracic source at T7 (275mm below the brainstem; Figure 3), we selected this as it minimises edge artefacts seen at the cervical and lumbar extremities of the cord and it facilitates visual comparisons across models.

**Figure 3.**
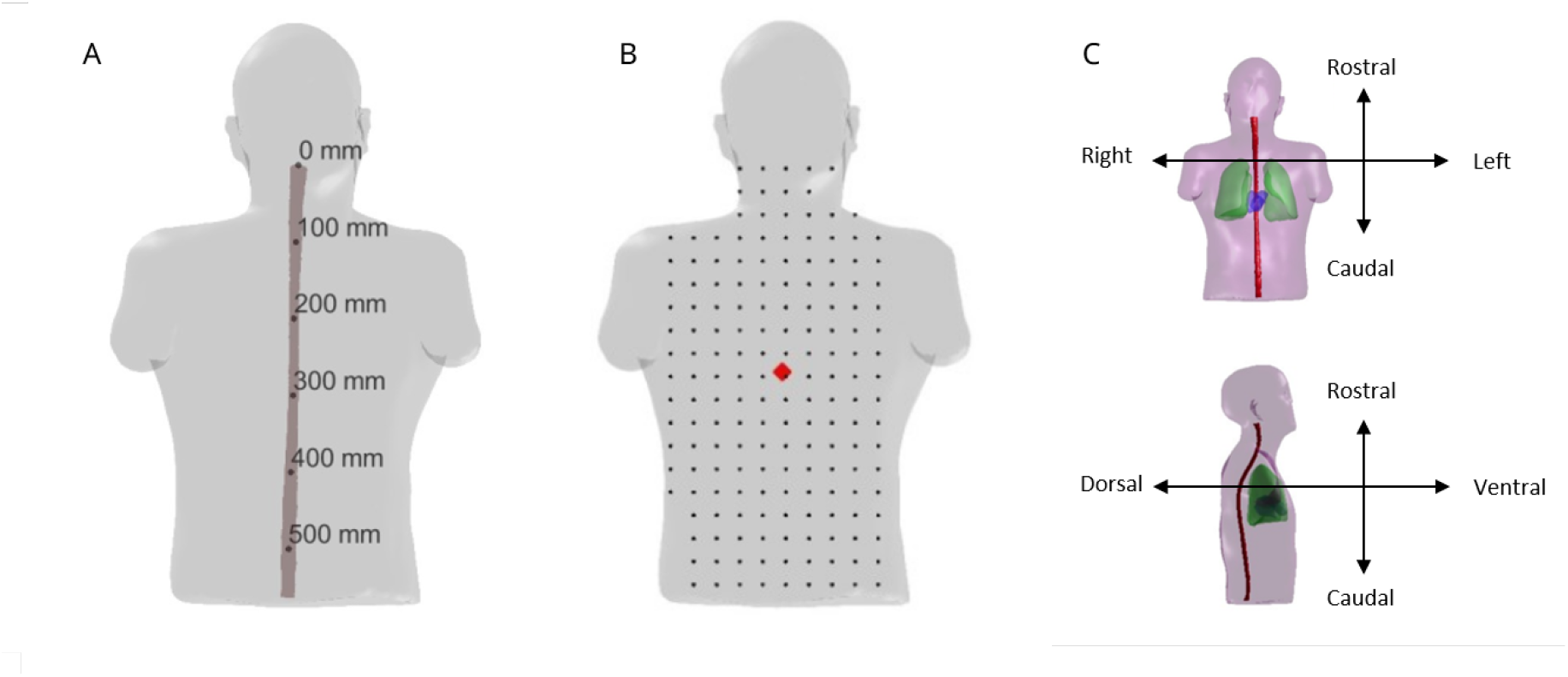
Anatomical context of spinal cord sources used for lead field analysis. **A:** source positions distributed along the spinal cord centreline at 5mm intervals, shown as distance from the brainstem (mm). **B:** posterior sensor array (dots) and the representative T7 reference source (red diamond) used for topographic visualisations throughout. **C:** Anatomical references for directions discussed; rostral–caudal, left–right and ventral–dorsal

Homogeneous and inhomogeneous toroidal bone models produced negligible differences in predicted lead fields (Supplementary Note S2), with relative errors below 5% and *r*^2^ exceeding 0.99 across all source positions and both frameworks. These models are therefore referred to collectively as the *toroidal bone model* throughout, with results being displayed from the inhomogeneous variant.

Figure 4 presents radial sensor-level topographies for T7 across all three dipole orientations. Spatial field patterns were consistent across bone models, with only modest amplitude differences. For RC-oriented dipoles, toroidal and realistic geometries produced amplitudes approximately 17% higher than the continuous model (5.0 vs. 6.0 fT/nAm), while spatial distributions were near-identical, with *r*^2^ between bone models exceeding 0.99 for RC-oriented sources at T7. LR-oriented dipoles showed a progressive amplitude increase with anatomical realism; VD-oriented dipoles exhibited minimal amplitude variation, with subtle asymmetries in the realistic model.

**Figure 4.**
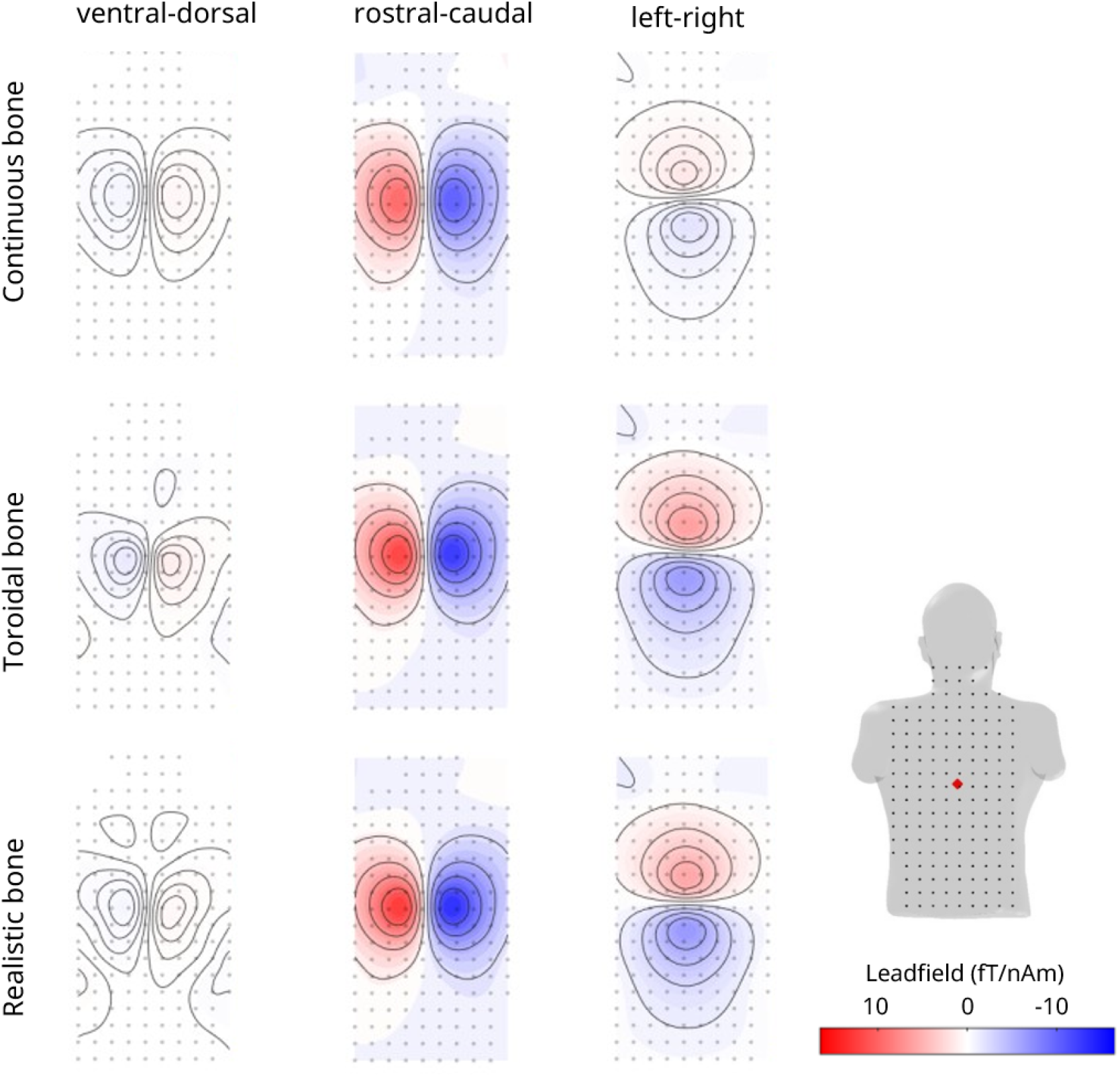
BEM radial sensor-level topographies for the representative T7 source using the posterior sensor array. The three rows correspond to the 3 bone models tested: continuous, toroidal and realistic. Columns correspond to dipole orientation (VD: ventral–dorsal; RC: rostral–caudal; LR: left–right); rows correspond to continuous, toroidal, and MRI-derived realistic bone models.

Across the full cord (Figure 5), all bone models followed consistent spatial sensitivity profiles. The dominant distinction was between continuous and segmented geometries. For LR-oriented sources, segmented models produced systematically higher amplitudes than the continuous model throughout the cord (35–67% for toroidal, 37–72% for realistic), with minimum values of 35% and 37% respectively confirming a cord-wide effect rather than a localised discrepancy. Segmented models also exhibited characteristic spatial modulation reflecting anatomical variation between vertebral segments and intervertebral regions. Note that despite relatively large modulation in magnitude with vertebral segment, there was only marginal (max modulation of (max modulation of *r*^2^ between continuous and realistic bone = 10% for the cervical region of the spinal cord) change in the correlation (i.e. the pattern) of the predicted field. Full quantitative pairwise comparisons are provided in Supplementary Tables S3 and S4.

**Figure 5.**
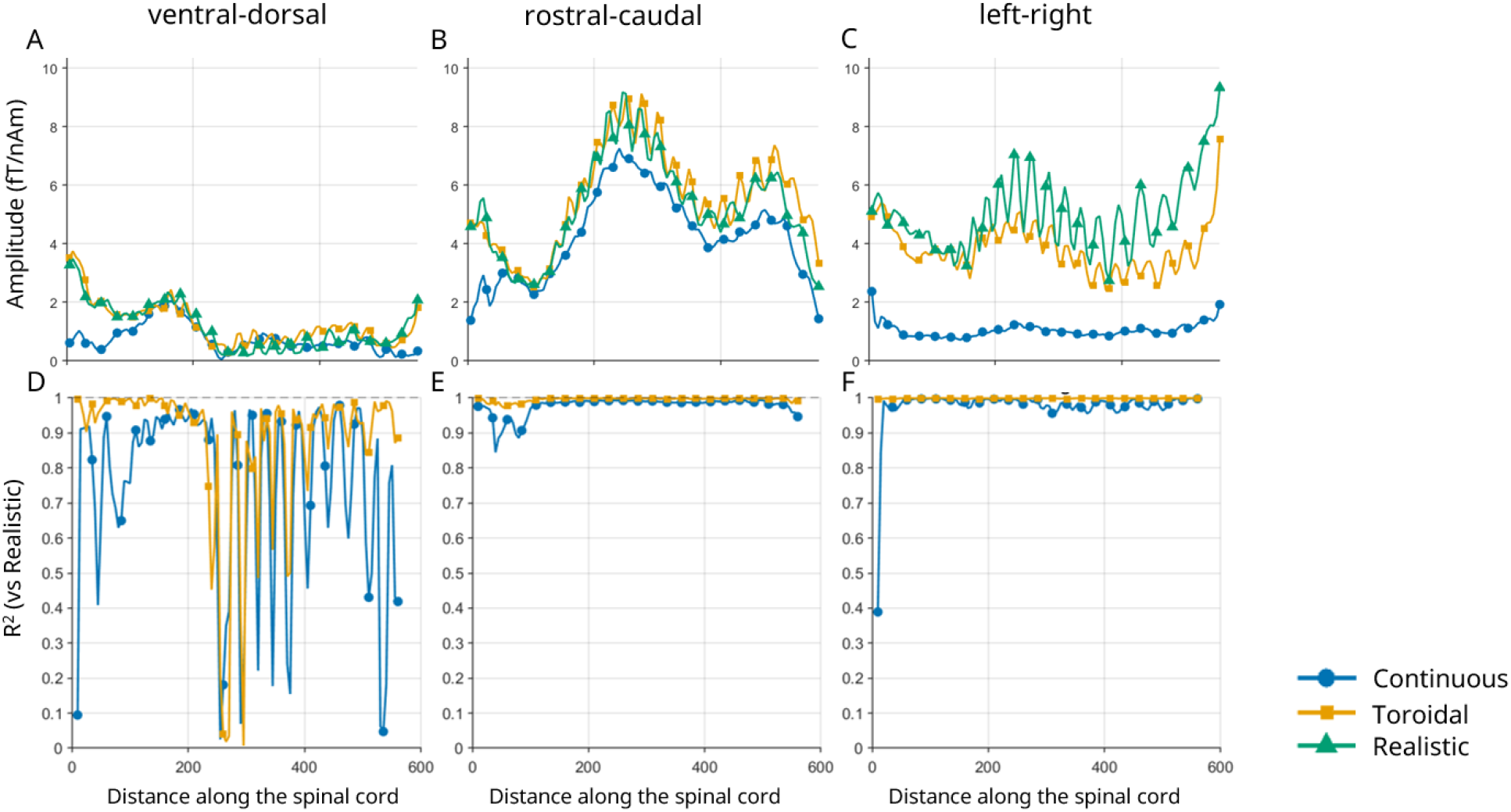
Lead field comparison across vertebral bone models using BEM. **(A–C)** Absolute maximum radial field amplitude (fT/nAm) for VD (ventral–dorsal; **A**), RC (rostral–caudal; **B**), and LR (left–right; **C**) dipole orientations. **(D–F)** Source-wise *r*^2^ relative to the MRI-derived realistic bone model for VD (**D**), RC (**E**), and LR (**F**) orientations. All panels show results across all cord positions for continuous (blue circles), toroidal (orange squares), and MRI-derived realistic (green triangles) bone models. The *x*-axis represents distance along the spinal cord from the brainstem (mm).

For RC- and LR-oriented sources, both the continuous and toroidal models maintained high *r*^2^ relative to the realistic bone model across the majority of the cord (*r*^2^ *>* 0.95), with the toroidal model showing particularly stable agreement even in the uppermost cord region (0–50mm) where the continuous model exhibited a modest reduction. For VD oriented sources, the continuous model showed pronounced source-wise volatility in *r*^2^, with values fluctuating between approximately 0.15 and over 0.8 between adjacent source positions along the full cord length. The toroidal model was considerably more stable for VD-oriented sources, maintaining *r*^2^ above 0.9 up to approximately 200mm, beyond which similarly pronounced variability emerged; this instability diminished progressively beyond 400mm. This volatility for VD sources reflects the near zero absolute amplitudes produced by this orientation in certain cord regions, where small absolute differences produce disproportionately large *r*^2^ fluctuations due to the leadfields being normalised by their amplitudes. This near-zero amplitude arises because, as shown in Figure 9, the VD dipole orientation becomes parallel to the outward torso surface normal in certain cord regions, placing these sources in a quasi-radial configuration in which the primary and secondary magnetic field contributions partially cancel, as described by Equation 7.

### Cross-Framework Comparison

Simulations performed using FEM yielded amplitude profiles and topographies closely mirroring those from BEM (Figure 6), with the same systematic distinction between continuous and segmented bone geometries present in both frameworks. Absolute maximum amplitude profiles and *r*^2^ relative to the realistic bone model across all FEM source positions were consistent with the BEM results reported above and are provided in Supplementary Note S3 (Figures S7–S8).

**Figure 6.**
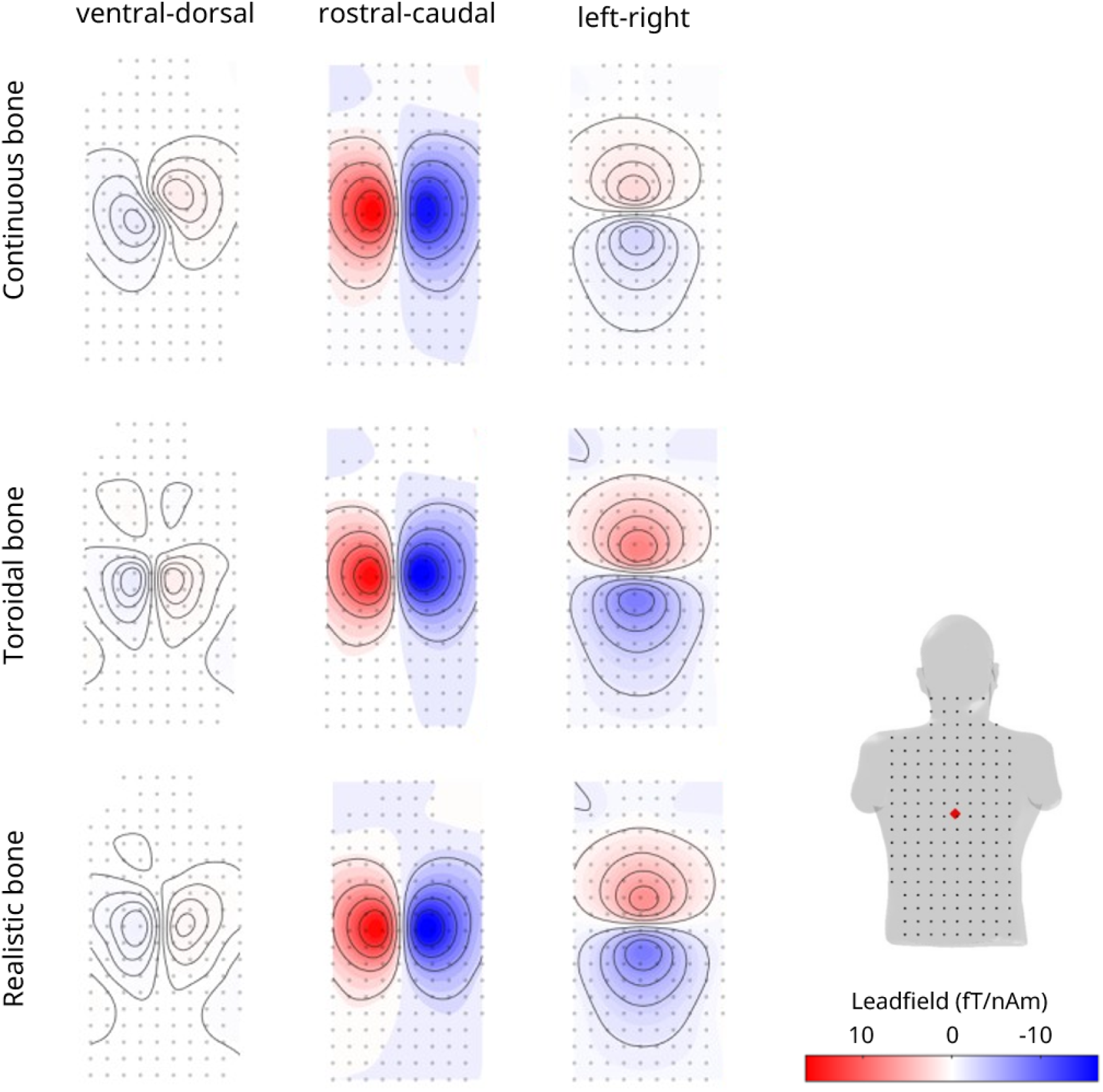
FEM radial sensor-level topographies for the representative T7 source using the posterior sensor array. Columns correspond to dipole orientation (VD: ventral–dorsal; RC: rostral–caudal; LR: left–right); rows correspond to continuous, toroidal, and MRI-derived realistic bone models. Format as in Figure 4.

Source-wise comparisons further investigate consistency between the BEM and FEM frameworks (Figure 7). For RC- and LR-oriented dipoles, *r*^2^ remained above 0.97 across all source locations and bone models, with RE below 15% throughout. For VD-oriented dipoles, *r*^2^ dropped as low as 0.40 in the 230–300mm thoracic region across all bone models, while RE remained below 21% across the same region. The most pronounced drop occurred at the source located 285mm along the cord, which was therefore selected for detailed topographic inspection.

**Figure 7.**
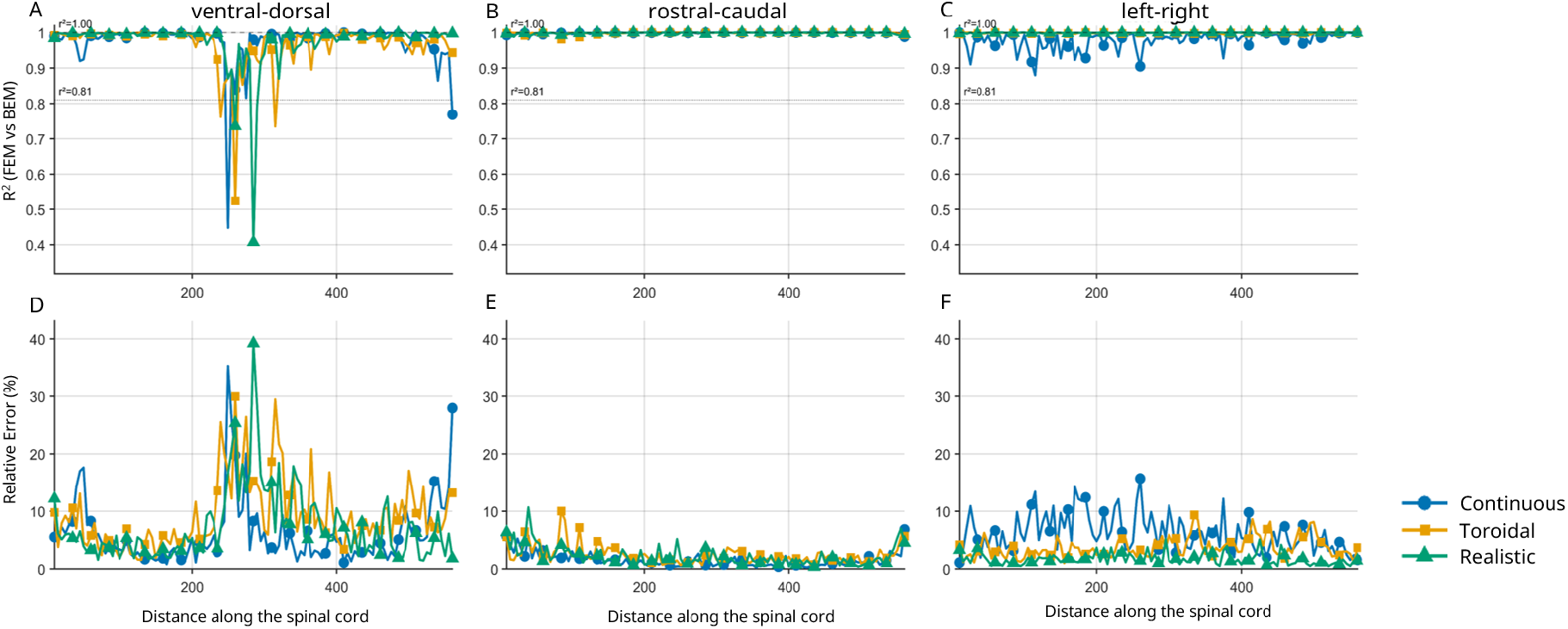
BEM–FEM framework comparison across all cord positions and bone models. **(a–c)** Source-wise *r*^2^ between BEM and FEM for VD (ventral–dorsal; **a**), RC (rostral–caudal; **b**), and LR (left–right; **c**) dipole orientations. **(d–f)** Relative error (RE) between BEM and FEM for VD (**d**), RC (**e**), and LR (**f**) orientations. All panels show results for continuous (blue circles), toroidal (orange squares), and MRI-derived realistic (green triangles) bone models across all cord positions. The *x*-axis represents distance along the spinal cord from the brainstem (mm).

Figure 8 shows the radial sensor-level topographies predicted by BEM and FEM at 285mm for the realistic bone model, using the VD dipole orientation at which the *r*^2^ drop is most severe. Despite producing comparable maximum amplitudes (approximately 0.6 fT/nAm), the two frameworks predict fundamentally different spatial field patterns at this source location: both produce dipolar responses, but the dipole axes are approximately perpendicular to one another.

**Figure 8.**
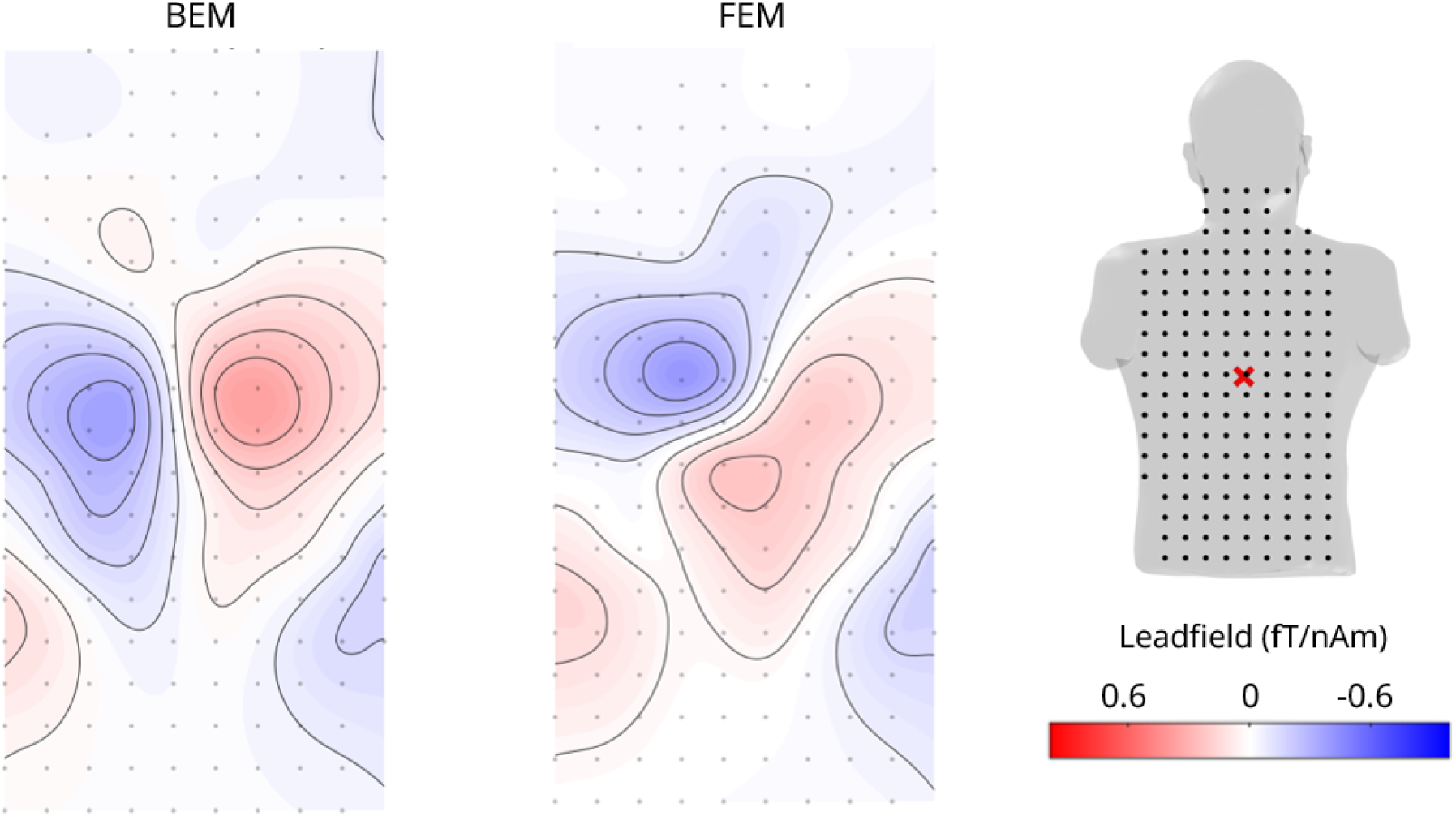
Radial sensor-level topographies at 285mm for the VD orientation using the MRI-derived realistic bone model. **Left:** BEM prediction. **Right:** FEM prediction. Despite comparable maximum amplitudes (≈ 0.6 fT/nAm), the two frameworks predict near-orthogonal field patterns: BEM produces positive and negative lobes oriented left–right, whilst FEM produces lobes oriented ventral–dorsal.

To investigate the geometric origin of this orientation-specific discrepancy, the angle between the outward surface normals of the torso and the orientation of each source dipole was computed along the full cord length. For each source, the outward normals of all torso surface vertices within a radius of 40mm (twice the sensor spacing) of the closest posterior sensor were averaged, and the angle between this averaged normal and the source dipole orientation was calculated. Figure 9 shows this angle as a function of source position along the cord for all three source orientations, alongside a 2D illustration of how the angle between source orientation and averaged surface normal is calculated.

For VD-oriented sources, the angle between the dipole and the surface normal reaches a minimum of approximately 0° in the 260–290mm cord region, indicating that these sources are effectively radially oriented with respect to the posterior torso surface. The surrounding sources within the 230–300mm region subtend angles below 10°, confirming that this is a zone of “quasi-radial” VD sources rather than a single isolated point. This geometric condition is the condition required for which the external magnetic field is generated exclusively by secondary volume currents, with no primary source contribution, as discussed in the context of radial silence in MEG^31, 34^. The perpendicular topographic disagreement between BEM and FEM observed at 285mm (Figure 8) is therefore a direct consequence of this near-radial geometry: when the primary field contribution vanishes, the predicted field is determined entirely by secondary currents at conductivity boundaries, and any difference in how those boundaries are represented between the two frameworks produces a fundamentally different predicted field orientation rather than a simple amplitude scaling error.

**Figure 9.**
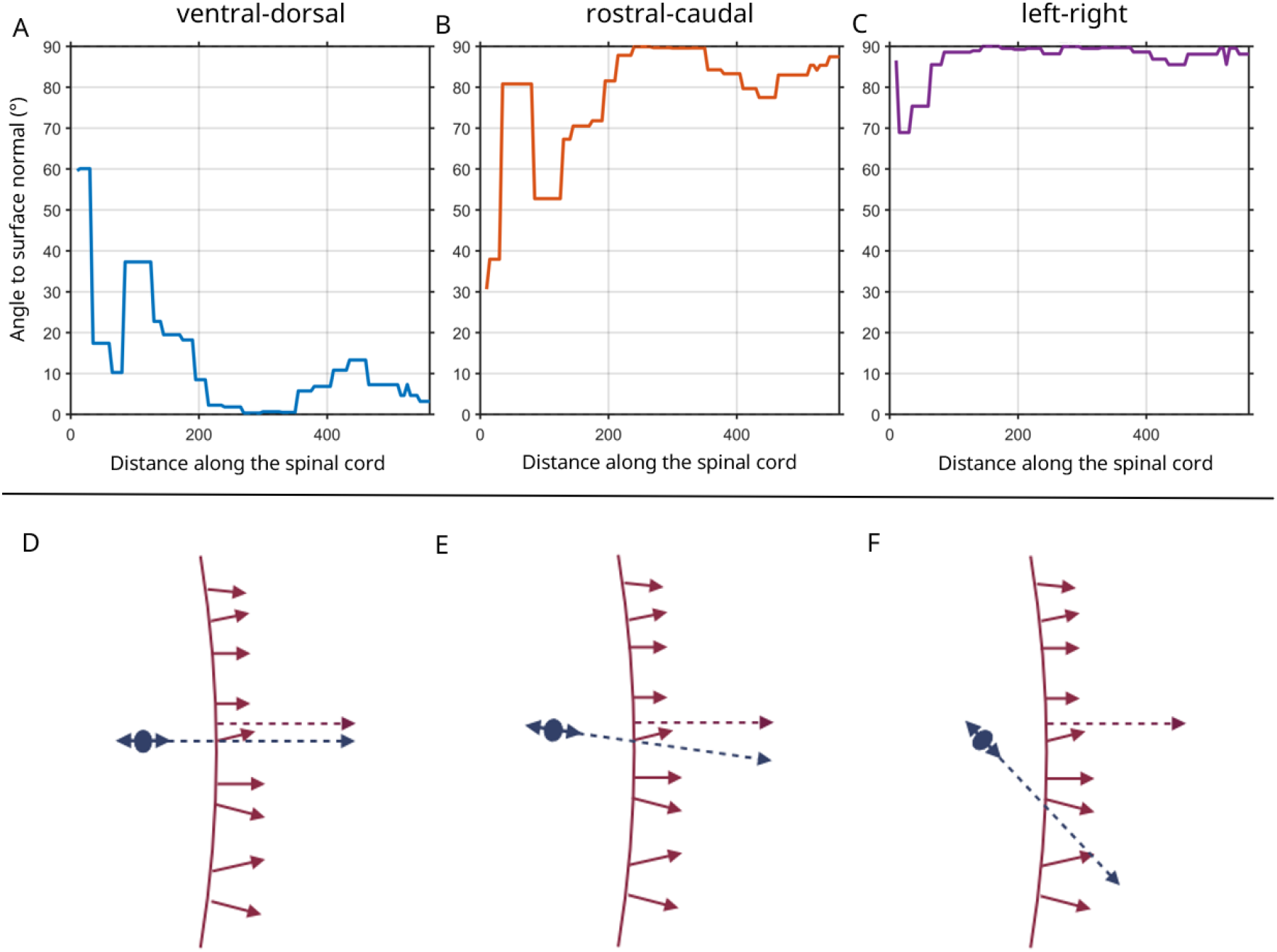
Surface normal analysis of source dipole geometry relative to the posterior torso surface. **(a-c)** Angle between the averaged outward torso surface normal and the source dipole orientation as a function of source position along the spinal cord (mm from brainstem), shown separately for VD (ventral–dorsal) (A), RC (rostral–caudal) (B), and LR (left–right) (C) orientations. An angle of 0° indicates a perfectly radial source; 90° indicates a perfectly tangential source. VD-oriented sources approach 0° in the 260–290mm region, confirming quasi-radial geometry. **(D-E)** Schematic of the geometric relationship between dipole orientation and averaged surface normal for perfectly radial **(D)**, quasi-radial **(E)**, and tangential **(F)** configurations.

In contrast, RC-oriented sources maintained angles above 65° across the majority of the cord, with only a modest reduction near the cervical region, whilst LR-oriented sources showed angles above 70° throughout the full cord length. This consistent near tangential geometry for RC and LR sources explains their uniformly high *r*^2^ across all cord positions: a large primary source contribution dominates the external field, reducing the relative influence of secondary boundary current differences between BEM and FEM.

To isolate the contribution of the heart and lungs to the BEM-FEM discrepancy at these quasi-radial source locations, BEM lead fields were recomputed with the heart removed, the lungs removed, and both removed simultaneously, and compared against the BEM original and FEM original. Full results are provided in Supplementary Note S3 (Figures S12–S17; Tables S5– S6). Within-BEM comparisons confirmed that organ removal produced only modest perturbations to the lead field: across all sensor axes, *r*^2^ between the BEM original and all organ-removal variants exceeded 0.969 for VD-oriented sources (Table S3), and RC- and LR-oriented sources showed median RE below 4.5% and *r*^2^ above 0.993 throughout. Critically, the localised BEM–FEM discrepancy for VD-oriented sources in the quasi-radial zone was not resolved by organ removal, removing both organs simultaneously worsened it, increasing the matched-pair median RE from 21.4% to 27.1% (Tables S5 and S6) and still producing the variability (in *r*^2^ and RE) between adjacent source positions as observed in Figure 7 at 200-400mm along the cord.

BEM–FEM agreement was verified for both tangential sensor axes, with matched pairs achieving median RE below 6.2% and median *r*^2^ above 0.981, consistent with the radial results (Table S1; Supplementary Section S1).

### Sensor Array Placement

Simulations were repeated using an anterior sensor array, across the chest, to assess the influence of sensor placement. BEM and FEM predictions were consistent for the anterior configuration, with spatial topographies and amplitude profiles closely matching the posterior results for equivalent bone models (Figure 10).

**Figure 10.**
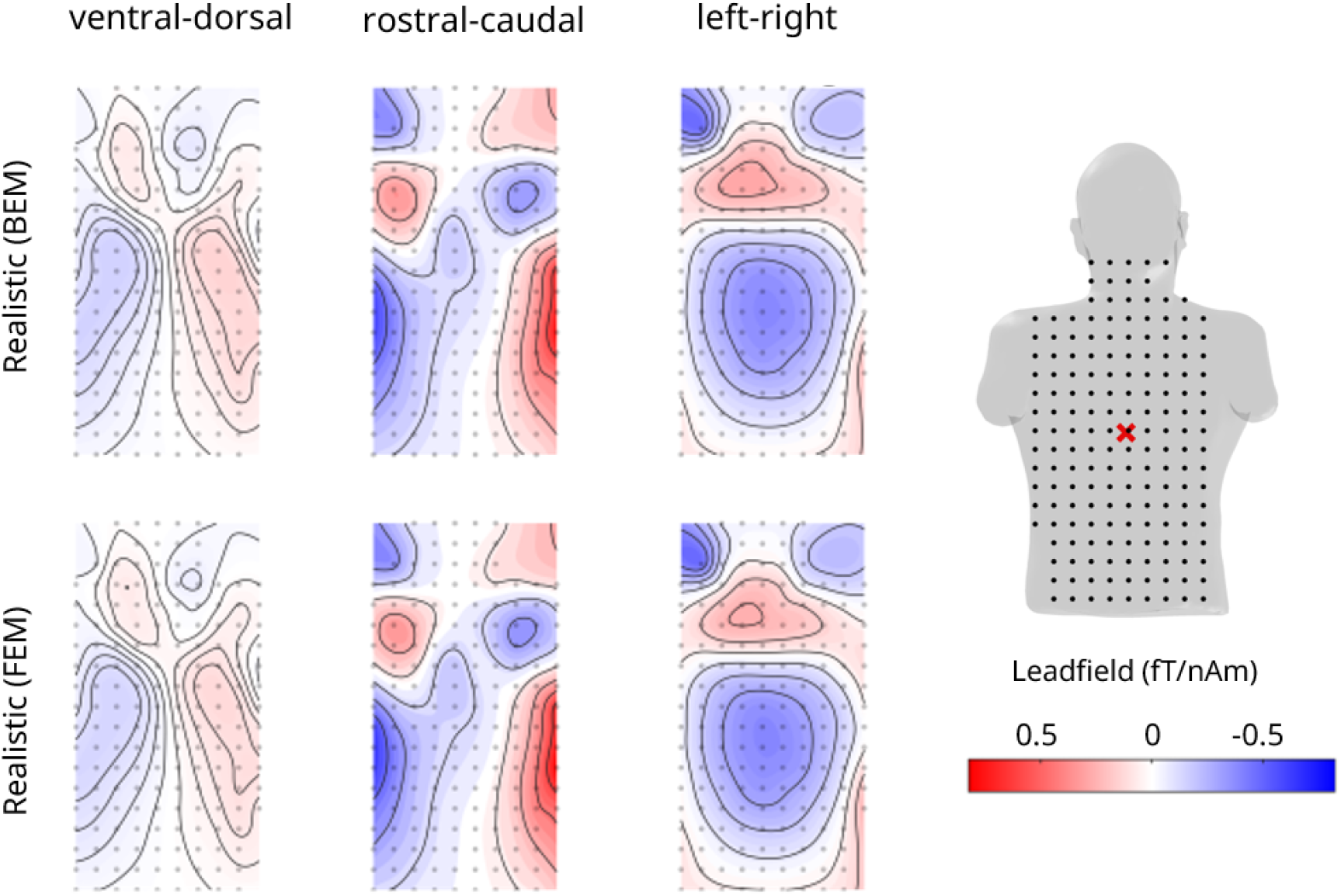
Anterior array radial sensor-level topographies for the representative T7 source. Columns correspond to dipole orientation (VD: ventral–dorsal; RC: rostral–caudal; LR: left–right); rows correspond to the realistic bone model for the BEM (top row) and FEM (bottom row).

Posterior arrays produced larger signal amplitudes overall, reflecting reduced source–sensor distance. However, anterior-to-posterior amplitude ratios (Figure 11) revealed that anterior sensitivity becomes comparable to posterior sensitivity in the cervical region, and this pattern was consistent between BEM and FEM. Framework-dependent differences were consistently smaller than geometry-dependent differences across all conditions.

**Figure 11.**
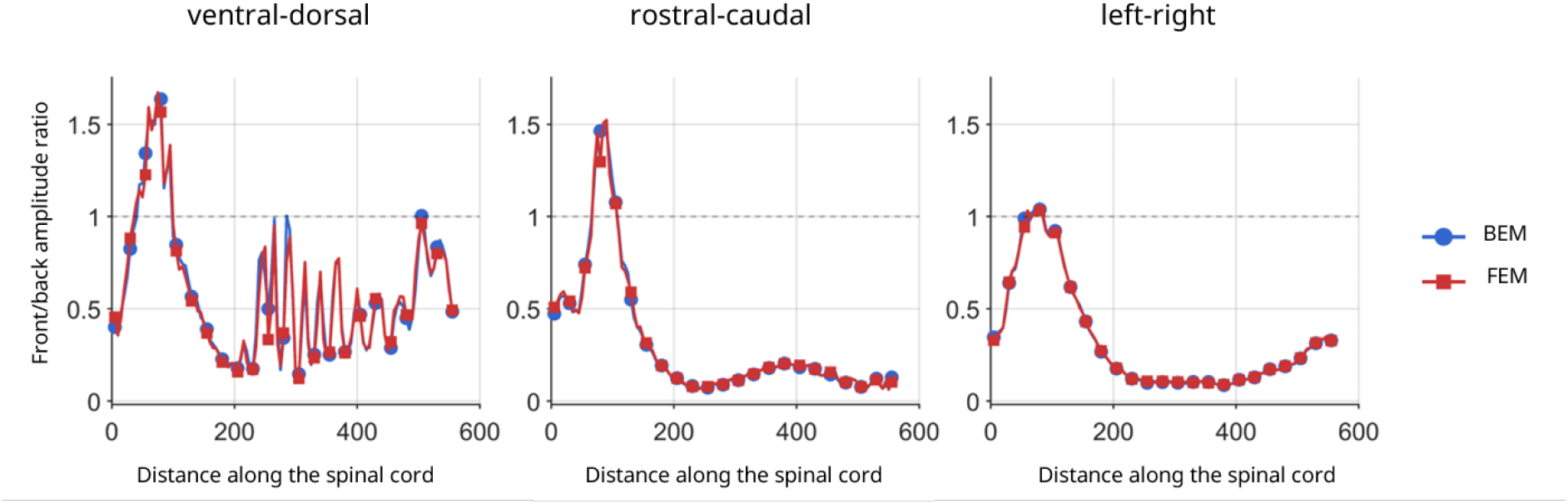
Anterior-to-posterior peak lead field amplitude ratio as a function of source position along the spinal cord. Ratios are shown for BEM (blue circles) and FEM (red squares) separately, across all dipole orientations (VD: ventral–dorsal; RC: rostral–caudal; LR: left–right) and bone models. A ratio of 1.0 indicates equal sensitivity; values below 1.0 indicate greater posterior sensitivity. The *x*-axis represents distance along the spinal cord from the brainstem (mm).

## Discussion

This study investigated how vertebral bone geometry, forward modelling framework, and sensor placement influence predicted MSG lead fields. The central finding is that BEM and FEM produce consistent forward solutions when equivalent anatomical geometries and conductivity assignments are used, and that the dominant source of variation in predicted lead fields is vertebral bone representation rather than solver choice.

### Influence of Bone Geometry

Vertebral bone geometry emerged as the most influential anatomical factor in MSG forward modelling. The dominant distinction was between the continuous bone representation and models incorporating vertebral segmentation, rather than between simplified and anatomically detailed segmented models.

For the continuous model, all three dipole orientations produced smooth dipolar topographies consistent with the right-hand rule, with RC sources generating the largest amplitudes and VD sources the smallest. Segmented geometries preserved these orientation dependent patterns but altered spatial field distributions, most notably producing spatially extended, less uniform VD topographies (Figures 4 and 6).

Anatomically-dependent effects were most pronounced for LR-oriented sources, where segmented geometries produced systematically higher amplitudes than the continuous model across the full cord (35–67% toroidal, 37–72% realistic). The consistency of these differences across all source positions indicates a cord-wide modulation of lateral current pathways rather than a localised boundary effect. This reflects the alignment between LR dipole orientation and the primary current paths modulated by vertebral segmentation: LR currents traverse bone laterally and are therefore most sensitive to changes in geometric representation, while VD currents are directed largely perpendicular to the sensor array and produce weaker, diffuse fields in all models (Figure 5).

Crucially, simplified toroidal and realistic MRI-derived bone models produced similar lead field amplitudes and spatial patterns across both frameworks, with differences between segmented geometries remaining well below those separating either from the continuous model (BEM: median RE 9.0%, FEM: median RE 11.5%; Supplementary Tables S3–S4). This demonstrates that the main geometric effect arises from representing vertebral segmentation itself rather than from fine-scale anatomical detail, and that simplified toroidal models provide a computationally efficient and physically adequate approximation to realistic vertebral anatomy for MSG forward modelling.

### Comparison Between BEM and FEM

A central objective was to evaluate whether the surface-based BEM formulation remains valid for the complex anatomy of the spinal cord and vertebral column. Despite its conceptual reduction of a volumetric problem to surface integrals over tissue boundaries, BEM produced forward solutions that were consistent with FEM across all bone models. Matched-pair median RE remained below 3.1% and median *r*^2^ exceeded 0.998, with the best agreement achieved for the most anatomically complex model (BEM Realistic vs. FEM Realistic: median RE 1.9%, median *r*^2^ 0.999, minimum *r*^2^ 0.992; Supplementary Table S4). Theoretically, the external magnetic field is determined by the net current crossing conductivity interfaces rather than by detailed internal current distributions. BEM captures this directly through its surface integrals, and the volumetric current resolution of FEM, which is where it differs from BEM, provides little advantage for the external field predictions we evaluated here. These findings are consistent with prior BEM–FEM validation in MEG^33, 48^.

The localised reduction in *r*^2^ for VD-oriented sources in the 230–300mm thoracic region warrants mechanistic consideration. As demonstrated by the surface normal analysis (Figures 8 and 9), VD-oriented sources in the 260–290mm region are effectively radially oriented with respect to the posterior torso surface, with the angle between the dipole axis and the averaged outward surface normal approaching 0°. The surrounding sources within the 230–300mm zone subtend angles below 10°, confirming this as a region of “quasi-radial” VD sources. In a spherically symmetric conductor, such a configuration produces no external magnetic field, because the cross-product term in Sarvas’s formula vanishes for a radially directed dipole^31, 34^. In the present geometry the torso is not spherically symmetric and the VD dipole is not perfectly radial, but the physical consequence is the same in principle: the primary current contribution to the external field is minimised and the predicted field at the posterior sensor array is generated almost exclusively by secondary volume currents at conductivity boundaries^29, 49^. This is precisely the condition under which the two numerical frameworks diverge most strongly.

The fundamental reason for this divergence lies in how BEM and FEM compute secondary volume currents near the outer mesh boundary under near-radial conditions. FEM solves for the electric potential throughout the full three-dimensional volume, allowing direct and continuous resolution of volume currents even in regions of steep potential gradient adjacent to conductivity boundaries^50^. Such gradients arise when a source is oriented to a nearby tissue interface, as the primary current drives charge accumulation at the boundary, causing a steep spatial transition in the electric potential which must be solved via the secondary magnetic fields to form a precise computation. The near cancellation between primary and secondary magnetic fields required for a quasi-radial source is therefore preserved within the FEM solution. BEM, by contrast, represents the potential only on two-dimensional surfaces; when a quasi-radial source produces an extreme potential gradient close to a surface boundary, the discrete surface elements approximate this gradient less accurately, which in turn affects the computed secondary current contribution to the magnetic field^36^. The result is not that one framework is correct and the other wrong, both are valid numerical approximations of the same underlying physics, but rather that the two methods resolve the secondary current structure differently under this specific geometric condition, producing predicted field topographies that diverge from one another. This divergence is what is observed as the near-orthogonal field orientations in Figure 8, both frameworks produce physically plausible dipolar responses, but the secondary current distributions they compute differ sufficiently that the resulting topographies point in different directions. The effect is further compounded by the presence of multiple high-contrast tissue boundaries in the thoracic region, the high-conductivity cardiac blood masses (0.62 S/m) and the low-conductivity lungs (0.05 S/m), which increase the complexity of the secondary current pathways that must be accurately resolved for the near-cancellation to be preserved^51^.

A direct practical consequence of the quasi-radial geometry is that absolute VD amplitudes in this region are very small, below 0.8 fT/nAm at the worst-case comparison sources, compared to RC amplitudes exceeding 9 fT/nAm at the same locations (Supplementary Table S3). When the true signal approaches zero, any difference in how secondary currents are resolved between the two frameworks produces a disproportionately large effect on both RE and *r*^2^, since both metrics are normalised by signal magnitude.

The original hypothesis for this divergence was the presence of the different conductivity boundaries in the chest cavity (caused by the heart and lungs) at approximately 200-400mm along the cord. However organ-removal analysis (Supplementary Figures S12–S17; Tables S5–S6) further confirms that the divergence is not driven by a simple difference in conductivity assignment. The localised BEM–FEM discrepancy for VD-oriented sources was not resolved by removing either organ, and removing both organs simultaneously worsened it, increasing the matched-pair median RE from 21.4% to 27.1% and producing still producing *r*^2^ oscillations between 0.375 and 1.0 at adjacent source positions. This is consistent with the secondary current interpretation: the divergence arises from how the two frameworks represent the geometrically complex organ boundaries that govern secondary current pathways in the quasi-radial zone, rather than from the conductivity values assigned to those organs. Source-to-sensor distance analysis (Supplementary Figures S10 –S11) confirms that the divergence cannot be attributed to proximity effects, sources in the 230–300mm range have source-to-sensor distances comparable to adjacent cord regions where BEM–FEM agreement is high.

For the RC- and LR-oriented sources most relevant to practical MSG, BEM and FEM remain in close agreement across all cord positions and bone models. From a practical perspective, and given isotropic conductivity (see below), BEM offers substantially lower computational cost and reduced mesh complexity compared to FEM while maintaining equivalent accuracy for the source positions and orientations relevant to MSG.

### Sensor Placement and Practical Implications

In the cervical region, anterior sensor sensitivity approaches or matches that of posterior arrays. This has direct experimental relevance: to date, virtually all MSG recordings in humans have employed posterior-only sensor or electrode configurations ^17, 26, 27, 39^. The cervical spinal cord receives afferent input from peripheral nerves responsible for upper limb movement, making it of particular interest for somatosensory studies. Our results suggest that anterior cervical sensor placement could substantially expand spatial coverage and sensitivity in spinal cord electrophysiology, and that this advantage is equivalent under both BEM and FEM. This motivates systematic experimental investigation of combined anterior–posterior sensor arrays in future MSG studies.

The consistency between frameworks across both array configurations confirms that BEM-based sensitivity maps can be used reliably to guide sensor placement optimisation, including prospective evaluation of novel anterior configurations.

### Limitations and Future Directions

Several limitations of the present study warrant further investigation.

#### Isotropic conductivity

All simulations assumed isotropic tissue conductivities. Spinal white matter exhibits substantial anisotropy due to aligned ascending and descending fibre tracts, and incorporating MRI-derived conductivity tensors into FEM simulations would allow more realistic modelling of directional current flow. Prior MEG studies suggest anisotropy may alter lead field amplitudes by approximately 10–30%, particularly for sources aligned with fibre tracts^36, 52^. Given that LR-oriented sources showed the greatest sensitivity to bone geometry in the present study, anisotropy effects may further modulate this orientation preferentially.

#### Static anatomical geometry

The spinal cord undergoes physiological motion due to cardiac pulsation and respiration^53^. Although these displacements are small relative to typical source–sensor distances, they introduce time-varying perturbations to the forward model that are not captured by the static geometry assumed here. Dynamic forward models incorporating measured cord motion could improve accuracy for high-precision source localisation, particularly in the 230–300mm thoracic region where BEM–FEM agreement was most sensitive to small amplitude differences.

#### Experimental validation

As a simulation study, direct experimental validation of forward model predictions against empirical recordings remains an important next step, and will be essential for confirming whether simplified geometric models, particularly the toroidal bone approximation, provide sufficient accuracy for practical MSG applications.

#### Additional anatomical structures

The present models do not include ribs, sternum, or detailed vertebral processes, which may locally influence conductivity pathways, particularly in thoracic regions where LR amplitude differences between bone models were most pronounced. Future work incorporating these features may further refine forward model accuracy.

## Conclusions

BEM and FEM produce equivalent forward solutions for MSG when applied to matched anatomical geometries, with median relative errors below 3.1% and median *r*^2^ exceeding 0.998 across all bone models and sensor configurations.

Vertebral bone geometry was the dominant source of variation in predicted lead fields. Segmented representations produced substantially larger effects than solver choice, while toroidal and MRI-derived realistic models performed comparably, demonstrating that simplified segmented geometries are sufficient for most MSG forward modelling applications.

Anterior sensor arrays achieve sensitivity comparable to posterior arrays in the cervical region, motivating investigation of combined anterior–posterior configurations in future MSG and ESG studies.

The `msg-coreg` pipeline (https://github.com/maikeschmidt/msg_coreg) provides an open-source platform for participant-specific forward modelling applicable to OPM-based MSG ^27^, cryogenic SQUID-based MSG^17^, and electrospinography^39^, providing methodological foundations for reproducible source modelling in spinal cord electrophysiology.

## Acknowledgments

The authors would like to thank Barbara Dymerska for her work and contribution to setting up the bespoke MRI protocol described in this study.

## Author Contributions

M.S.: Conceptualisation, Methodology, Software, Validation, Formal Analysis, Investigation, Data Curation, Writing - Original Draft.

G.O’N.: Methodology, Software (FEM implementation), Writing - Review, Editing. M.E.S.: Conceptualization, Writing - Review, Editing.

S.M.: Methodology, Writing - Review, Editing.

S.B.: Resources, Writing - Review, Editing, Supervision.

M.F.C.: Conceptualisation, Resources, Writing - Review, Editing, Supervision, Funding Acquisition. G.R.B.: Conceptualisation, Resources, Writing - Review, Editing, Supervision, Funding Acquisition.

## Additional information

### Data and Code Availability

All anatomical processing and sensor array generation code is available at https://github.com/maikeschmidt/msg_coreg(release v1.0).

All forward modelling code (BEM and FEM) used in this study, as well as all analysis, is available at https://github.com/maikeschmidt/msg_fwd

Forward modelling foundations code for BEM (Helsinki BEM Framework) is available at https://github.com/MattiStenroos/hbf_lc_p

Forward modelling foundations code for FEM (DUNEuro + MATLAB interface) is available at https://github.com/georgeoneill/study-spinevol/tree/main

Processed meshes and lead field matrices are available upon reasonable request to the corresponding author. Raw MRI data cannot be shared due to participant privacy, but the processing pipeline enables generation of equivalent models from users’ own data.

