## supplementary materials for "Forward Modelling for Magnetospinography: Systematic Comparison of Boundary Element and Finite Element Methods"

### Supplementary Material

#### S1 BEM–FEM Comparison Across Tangential Sensor Axes

##### Overview

The main text reports BEM–FEM comparisons for the radial sensor axis (axis 3), which is the primary axis of interest for MSG as it captures the dominant field component from spinal cord sources measured at the posterior sensor array. To confirm that framework agreement is not specific to this sensor orientation, the same pairwise comparisons were performed for the two tangential axes (axis 1 and axis 2).

##### Sensor-level topographies

Supplementary Figures S1 and S2 show sensor-level topographies for T7 across all three sensor axes and all three dipole orientations for the BEM and FEM frameworks respectively. Across both frameworks, the spatial field patterns on the tangential axes are visually consistent with the radial axis results seen in the main text. The amplitude increase from continuous to segmented bone geometries observed on the radial axis is similarly present on both tangential axes. The anterior–posterior field distribution pattern, however, differs across the three bone models on the tangential axes, reflecting the directional sensitivity of these components to lateral and rostral–caudal current flow modified by vertebral geometry.

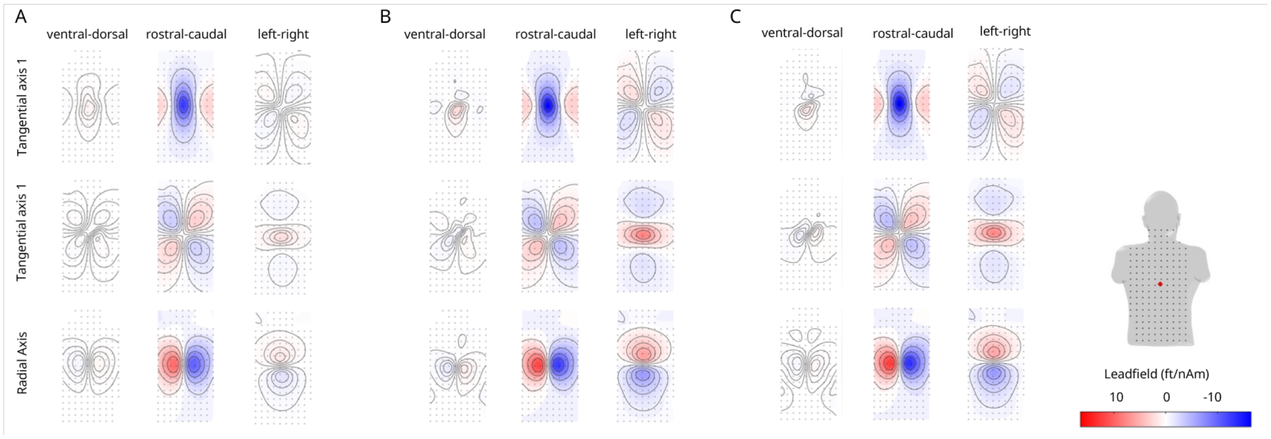


**Figure S1. Supplementary Figure S1 | BEM sensor-level topographies for T7 across all three sensor axes and dipole orientations.** Corresponding to continuous **(a)**, toroidal **(b)**, and MRI-derived realistic **(c)** bone models. Rows correspond to sensor axes 1, 2, and 3 (radial); within each subfigure, columns correspond to dipole orientations (VD: ventral–dorsal; RC: rostral–caudal; LR: left–right).

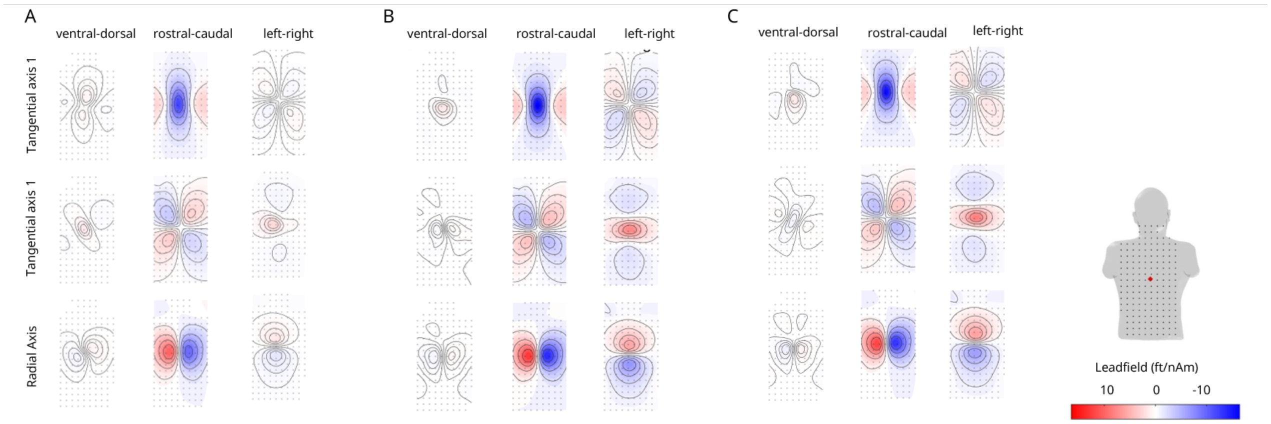


**Figure S2. Supplementary Figure S2 | FEM sensor-level topographies for T7 across all three sensor axes and dipole orientations.** Corresponding to continuous **(a)**, toroidal **(b)**, and MRI-derived realistic **(c)** bone models. Format as in Supplementary Figure S1.

##### Matched cross-framework agreement

Supplementary Table S1 summarises the matched BEM–FEM pair RE and $r^2$ metrics across all three sensor axes. On both tangential axes, matched pairs achieved median RE below 6.2% and median $r^2$ above 0.981 for all bone models, confirming that BEM–FEM agreement holds beyond the radial component. The toroidal and realistic matched pairs perform comparably on the tangential axes to the radial axis. BEM Continuous vs. FEM Continuous shows a modest increase in RE on axis 2 (6.1% vs. 2.5% on axis 3), reflecting the greater sensitivity of tangential components to internal volume current structure in the simpler continuous geometry, where current pathways are less constrained by vertebral boundaries. Segmented bone models show smaller axis 2 discrepancies (BEM Toroidal vs. FEM Toroidal: 4.7%; BEM Realistic vs. FEM Realistic: 3.5%), consistent with vertebral segmentation reducing the diffuse volume current contributions that primarily affect tangential components. Minimum $r^2$ values remain above 0.920 on axis 1 and above 0.890 on axis 2 for all matched pairs, confirming that spatial field patterns are well-preserved across frameworks on all sensor components.

**Table S1.** **Supplementary Table S1** | Matched framework pair (same bone model, BEM vs. FEM) median relative error (RE) and median squared correlation coefficient ($r^2$) across all three sensor axes. Axis 3 (radial) values are reproduced from the main text for comparison.

| | | Axis 1 (tangential) | | Axis 2 (tangential) | | Axis 3 (radial) | |
|---|---|---|---|---|---|---|---|
| **Model A** | **Model B** | **RE Med.** | **$r^2$ Med.** | **RE Med.** | **$r^2$ Med.** | **RE Med.** | **$r^2$ Med.** |
| BEM Continuous | FEM Continuous | 3.9% | 0.996 | 6.1% | 0.981 | 2.5% | 0.998 |
| BEM Toroidal | FEM Toroidal | 4.5% | 0.994 | 4.7% | 0.990 | 3.1% | 0.998 |
| BEM Realistic | FEM Realistic | 3.7% | 0.995 | 3.5% | 0.996 | 1.9% | 0.999 |

##### Geometry-dependent effects on tangential axes

The geometry-driven separation between continuous and segmented bone models observed on the radial axis is equally present on both tangential axes. Supplementary Table S2 reports minimum and maximum amplitude percentage differences for LR-oriented sources, which showed the most systematic bone geometry effects in the main text. On axis 1, BEM Continuous vs. BEM Toroidal differences range from 45.1–67.4% and BEM Continuous vs. BEM Realistic from 47.2–73.2%, with elevated minimum values confirming cord-wide systematic offsets. On axis 2, these ranges are 47.0–66.5% and 56.2–72.7% respectively, consistent with the radial axis values (35.0–66.6% and 36.6–72.4%) reported in Supplementary Table S3. The continuous bone model's systematic underestimation of LR-oriented signal amplitude is therefore not an artefact of any particular sensor component. The Toroidal–Realistic pair remained consistently tight across all tangential axes (axis 1: RE median 9.0%, $r^2$ median 0.980; axis 2: RE median 9.7%, $r^2$ median 0.980), mirroring the FEM within-framework equivalent (axis 1: 11.5%; axis 2: 12.9%), and confirming that the primary geometric distinction separates continuous from segmented bone models regardless of measurement axis.

**Table S2.** **Supplementary Table S2** | Minimum and maximum symmetric amplitude percentage differences for LR-oriented sources on the two tangential sensor axes, for the primary within-framework and matched cross-framework pairs. Matched framework pairs are shaded.

| | | Axis 1 | | Axis 2 | |
|---|---|---|---|---|---|
| **Model A** | **Model B** | Min (%) | Max (%) | Min (%) | Max (%) |
| *Within-framework comparisons (BEM)* | | | | | |
| BEM Continuous | BEM Toroidal | 45.1 | 67.4 | 47.0 | 66.5 |
| BEM Continuous | BEM Realistic | 47.2 | 73.2 | 56.2 | 72.7 |
| BEM Toroidal | BEM Realistic | 0.1 | 31.6 | 0.2 | 32.3 |
| *Cross-framework matched pairs (BEM vs. FEM)* | | | | | |
| BEM Continuous | FEM Continuous | 0.1 | 18.2 | 0.1 | 31.4 |
| BEM Toroidal | FEM Toroidal | 0.0 | 9.2 | 0.0 | 16.3 |
| BEM Realistic | FEM Realistic | 0.0 | 7.2 | 0.0 | 12.6 |

##### Axis-specific observations

The maximum amplitude percentage differences for RC- and VD-oriented sources on both tangential axes continue to occur at low-amplitude positions (below 0.2 fT/nAm), consistent with the radial axis behaviour and the caveat noted in Supplementary Table S3. On axis 1, worst-case VD differences consistently arise at the near-cord-surface positions (10 mm and 560 mm), driven by small absolute amplitude differences where the tangential component approaches zero. Taken together, the tangential axis results confirm that the core conclusions of the main text generalise across all measurable field components: BEM and FEM agree closely for matched bone models, the dominant geometric effect separates continuous from segmented bone, and the toroidal and realistic models behave similarly regardless of measurement axis.

#### S2 Homogeneous and Inhomogeneous Toroidal Bone Model Comparisons

##### Overview

Within the toroidal bone representation, two variants were considered: a homogeneous toroidal model in which each toroid has uniform geometry throughout the cord, and an inhomogeneous toroidal model in which toroid size and thickness vary anatomically along the spinal cord to approximate the gradual change in vertebral morphology. The main text collapses these into a single *toroidal* category on the basis of their close agreement. This section documents the quantitative basis for that decision.

##### BEM framework

Supplementary Figure S3 shows the absolute maximum lead field amplitude across all spinal cord source positions for the homogeneous and inhomogeneous toroidal BEM models. The two models are in close agreement throughout the cord for all dipole orientations, with a progressive divergence emerging at higher source positions (beyond approximately 400 mm). This divergence reflects the fact that the inhomogeneous model incorporates increasingly large and thick toroids toward the lower thoracic and lumbar levels, while the homogeneous model maintains constant geometry. Despite this anatomically motivated variation, the amplitude difference remains modest across the majority of the cord, supporting the use of the simpler homogeneous model as a representative of the toroidal class.

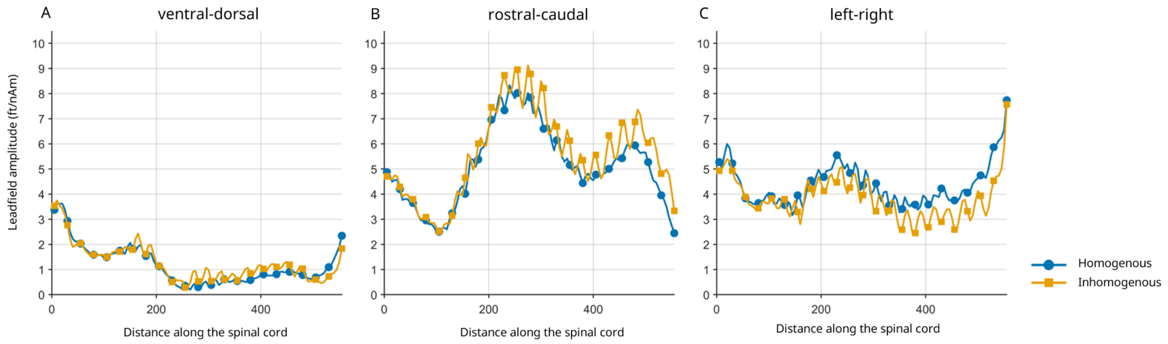


**Figure S3. Supplementary Figure S3 |** Absolute maximum lead field amplitudes (fT/nAm) for homogeneous (blue) and inhomogeneous (yellow) toroidal BEM bone models across all spinal cord source positions for VD (ventral–dorsal), RC (rostral–caudal), and LR (left–right) dipole orientations. The $x$-axis represents distance along the spinal cord from the brainstem (mm).

##### FEM framework

Supplementary Figure S4 shows the equivalent comparison within the FEM framework. The pattern mirrors the BEM result: homogeneous and inhomogeneous toroidal models are in close agreement along the majority of the cord, with the same gradual divergence toward higher source positions arising from anatomical variation in toroid geometry. The consistency of this pattern across frameworks further confirms that the small differences between toroidal variants are a genuine geometric effect rather than a framework-dependent numerical artefact.

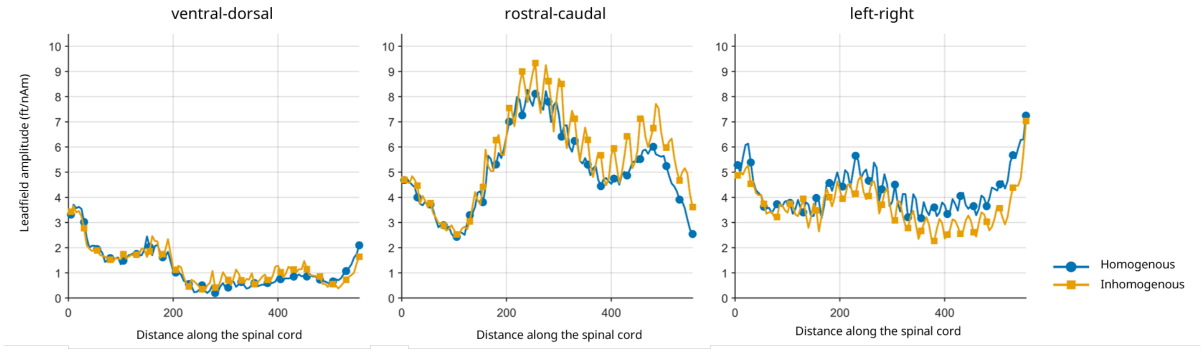


**Figure S4.** **Supplementary Figure S4 |** Absolute maximum lead field amplitudes (fT/nAm) for homogeneous and inhomogeneous toroidal FEM bone models across all spinal cord source positions. Format as in Supplementary Figure S3.

#### Cross-framework comparisons

Supplementary Figure S5 presents RE and $r^2$ metrics for the radial sensor axis comparing all four toroidal model combinations: homogeneous BEM (A), inhomogeneous BEM (B), homogeneous FEM (C), and inhomogeneous FEM (D). The matched cross-framework pairs achieved the closest agreement: homogeneous BEM vs. homogeneous FEM gave median RE of 2.3% and median $r^2$ of 0.9985, while inhomogeneous BEM vs. inhomogeneous FEM gave median RE of 3.0% and median $r^2$ of 0.9977. Within-framework comparisons between toroidal variants were slightly larger but remained small: homogeneous vs. inhomogeneous BEM gave median RE of 4.9% and $r^2$ of 0.9921, and homogeneous vs. inhomogeneous FEM gave median RE of 6.2% and $r^2$ of 0.9676. All four values are well within the range associated with the continuous-to-segmented bone transition reported in the main text, confirming that both toroidal variants belong to the same geometric class.

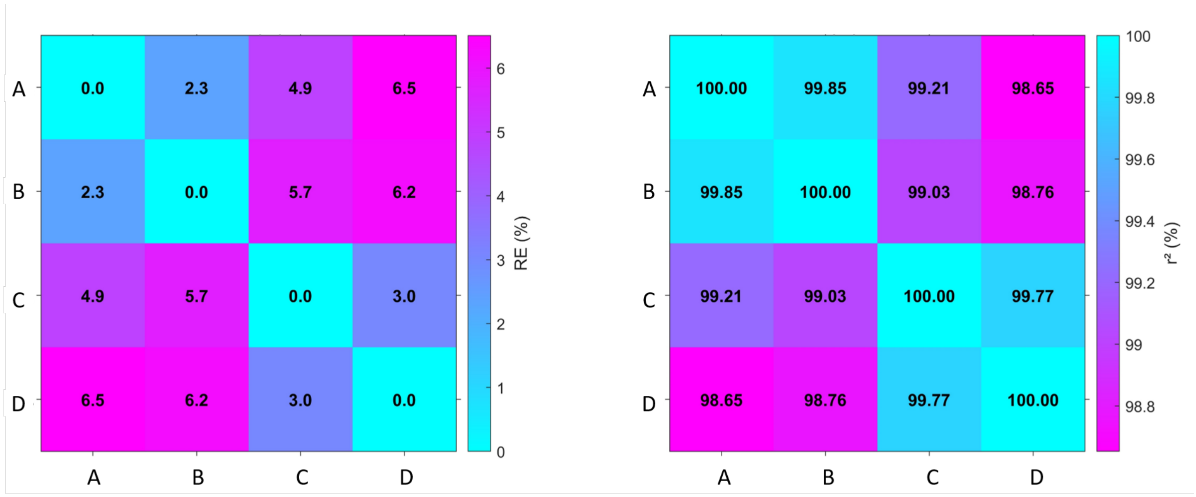


**Figure S5.** **Supplementary Figure S5 |** Relative error (RE) and squared correlation coefficient ($r^2$) for the radial sensor axis comparing homogeneous and inhomogeneous toroidal bone models across BEM and FEM frameworks averaged across all three dipole orientations (VD: ventral–dorsal; RC: rostral–caudal; LR: left–right). Labels: A = homogeneous BEM, B = inhomogeneous BEM, C = homogeneous FEM, D = inhomogeneous FEM.

Supplementary Figure S6 shows source-wise RE and $r^2$ for the matched cross-framework toroidal pairs. For LR-oriented sources, $r^2$ remained consistently above 0.995 across all source positions for both toroidal variants, with RE showing source-to-source variation reflecting the spatial modulation of vertebral anatomy, but not exceeding 9% for the inhomogeneous model and remaining below 6% for the homogeneous model throughout the cord. For RC-oriented sources, $r^2$ values were stable above 0.98 beyond approximately 180 mm, with the inhomogeneous model showing a slightly larger dip toward the near-cord-surface positions ($r^2$ minimum approximately 0.98 vs. 0.99 for homogeneous). For VD-oriented sources, both models showed the same pattern observed in the main text: stable $r^2$ above 0.95 outside the 200–300 mm thoracic region, with a localised drop to approximately 0.5 within this region, followed by gradual recovery.

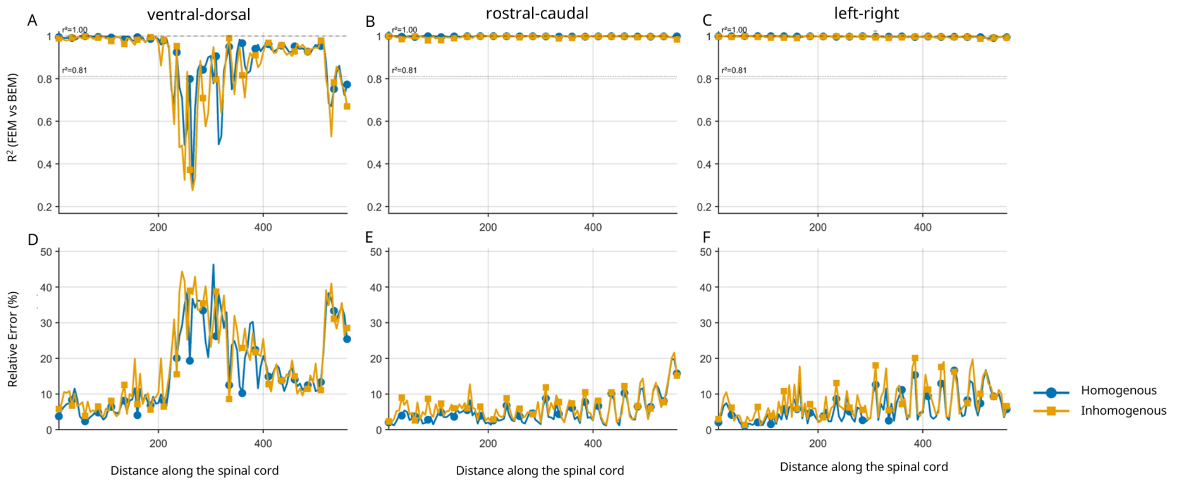


**Figure S6.** **Supplementary Figure S6 |** Source-wise squared correlation coefficient ($r^2$, top) and relative error (RE, bottom) for matched cross-framework toroidal pairs (homogeneous BEM vs. homogeneous FEM; inhomogeneous BEM vs. inhomogeneous FEM), shown separately for each dipole orientation (VD: ventral–dorsal; RC: rostral–caudal; LR: left–right). The $x$-axis represents distance along the spinal cord from the brainstem (mm).

### S3 Quantitative Cross-Framework and Bone Model Comparisons

#### Overview

This section provides full quantitative pairwise comparisons between all bone model and framework combinations using relative error (RE) and squared correlation coefficient ($r^2$), and includes additional figures and organ-removal analyses supporting the Results and Discussion. The main text reports summary findings for the radial sensor axis; full per-orientation and per-variant breakdowns are given here.

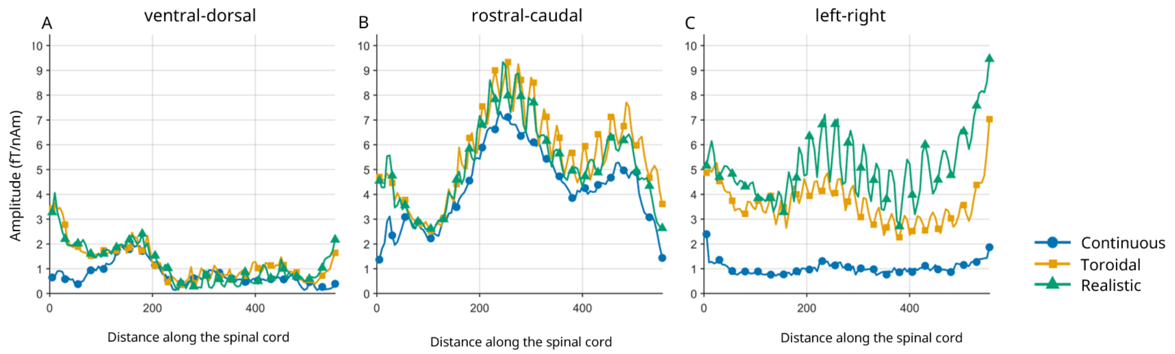


**Figure S7.** **Supplementary Figure S7 |** Absolute maximum lead field amplitudes (fT/nAm) across all spinal cord sources using FEM for continuous (blue), toroidal (orange), and MRI-derived realistic (green) bone models, shown separately for each dipole orientation (VD: ventral–dorsal; RC: rostral–caudal; LR: left–right). Results are consistent with the BEM equivalent reported in Figure 5(a–c) of the main text.

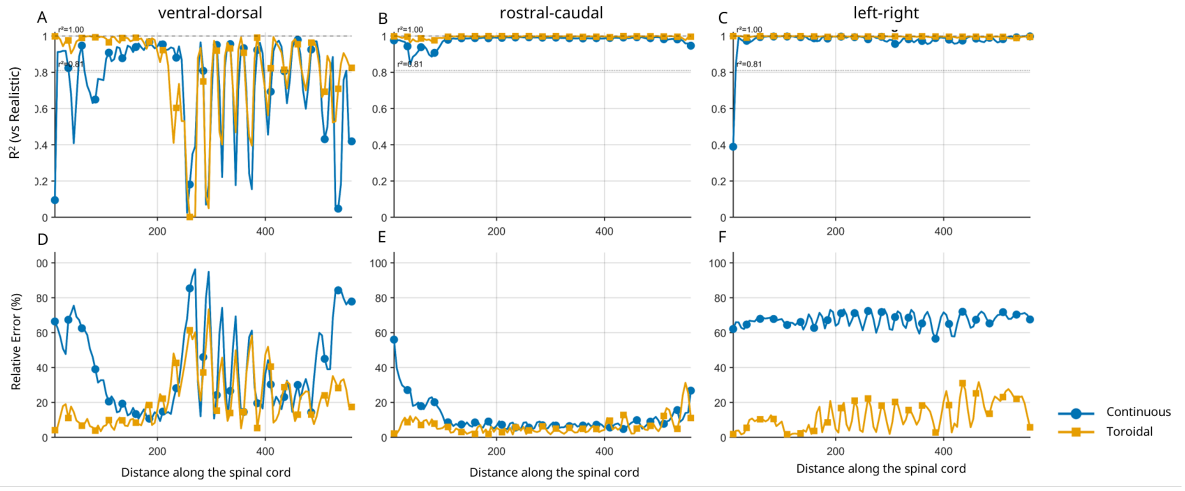


**Figure S8.** **Supplementary Figure S8 |** Source-wise $r^2$ of the continuous and toroidal bone models relative to the MRI-derived realistic bone model across all source positions using FEM, shown separately for each dipole orientation (VD: ventral–dorsal; RC: rostral–caudal; LR: left–right). Results mirror the BEM equivalent reported in Figure 5(d–f) of the main text.

#### Full pairwise BEM–FEM comparisons across all bone models

Supplementary Figure S9 and Tables S3–S4 provide complete pairwise RE and $r^2$ comparisons across all bone model and framework combinations for the radial sensor axis. These complement the organ-removal analyses above and the summary statistics reported in the main text. Note that in Supplementary Figure S9 we show both the homogenous and inhomogeneous toroidal bones, but as discussed above, the tables show results for only the inhomogeneous bone variant which has been labelled the 'toroidal' bone.

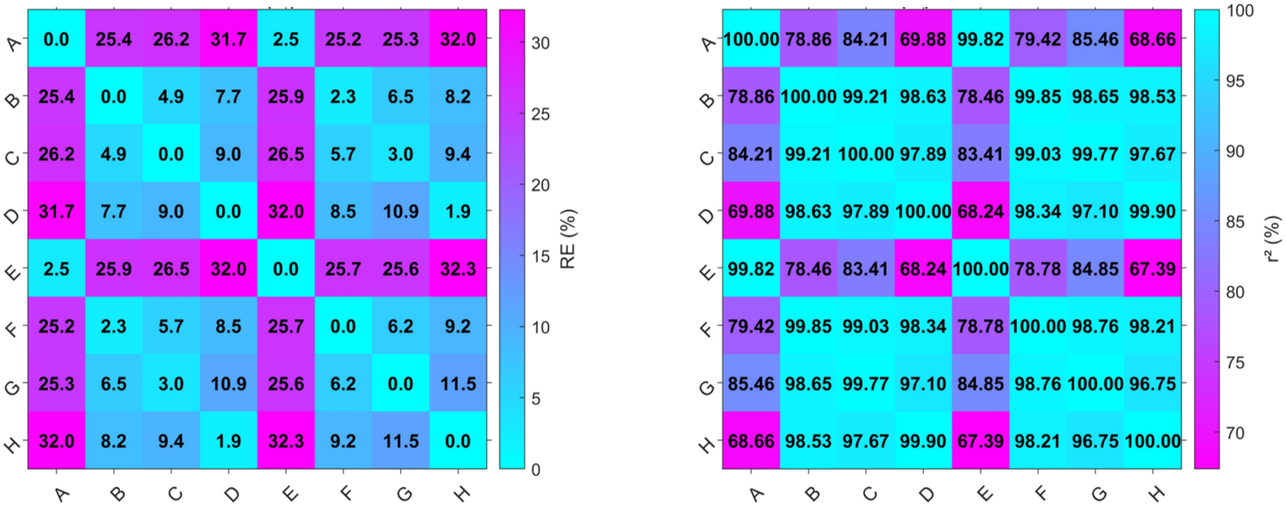


**Figure S9.** **Supplementary Figure S9 |** Relative error (RE) and $r^2$ comparing BEM and FEM lead fields for continuous, homogenous, inhomogeneous and realistic bone models across all pairwise combinations. Labels denote: A: BEM Continuous, B: BEM Homogenous, C: BEM Inhomogeneous, D: BEM Realistic, E: FEM Continuous, F: FEM Homogenous, G: Inhomogeneous, H: Realistic. Matched cross-framework pairs (A–E, B–F, C–G, D–H) are the primary comparison of interest.

**Table S3. Supplementary Table S3 | ** Minimum and maximum symmetric percentage amplitude differences between bone models across all spinal cord source locations, computed for the radial sensor axis. Values are computed as $|A-B|/(|A|+|B|)\times 100$. Matched framework pairs (same bone model, BEM vs. FEM) are shaded. Entries marked * have their maximum at a source position where absolute amplitude <0.2 fT/nAm; these large apparent differences are a consequence of near-zero VD signal in the mid-thoracic cord region rather than genuine forward model error.

| Model A | Model B | Ori. | Min (%) | Max (%) | Position (mm) |
|---|---|---|---|---|---|
| *Within-framework comparisons (BEM)* | | | | | |
| BEM Continuous | BEM Toroidal | LR | 35.0 | 66.6 | 180 |
| BEM Continuous | BEM Realistic | LR | 36.6 | 72.4 | 290 |
| BEM Toroidal | BEM Realistic | LR | 0.1 | 34.9 | 435 |
| BEM Continuous | BEM Toroidal | RC | 1.1 | 54.5* | 10 |
| BEM Continuous | BEM Realistic | RC | 0.4 | 53.6* | 10 |
| BEM Toroidal | BEM Realistic | RC | 0.1 | 25.9 | 550 |
| BEM Continuous | BEM Toroidal | VD | 0.0 | 83.3* | 250 |
| BEM Continuous | BEM Realistic | VD | 0.3 | 78.5* | 545 |
| BEM Toroidal | BEM Realistic | VD | 0.1 | 62.5* | 300 |
| *Cross-framework matched pairs (BEM vs. FEM)* | | | | | |
| BEM Continuous | FEM Continuous | LR | 0.0 | 16.4 | 300 |
| BEM Toroidal | FEM Toroidal | LR | 0.0 | 9.1 | 335 |
| BEM Realistic | FEM Realistic | LR | 0.0 | 5.2 | 295 |
| BEM Continuous | FEM Continuous | RC | 0.0 | 4.9 | 25 |
| BEM Toroidal | FEM Toroidal | RC | 0.0 | 4.0 | 560 |
| BEM Realistic | FEM Realistic | RC | 0.0 | 6.7 | 40 |
| BEM Continuous | FEM Continuous | VD | 0.0 | 45.4* | 250 |
| BEM Toroidal | FEM Toroidal | VD | 0.1 | 21.2* | 275 |
| BEM Realistic | FEM Realistic | VD | 0.0 | 17.1* | 340 |

**Table S4. Supplementary Table SS4 | ** Summary of pairwise relative error (RE) and squared correlation coefficient ($r^2$) across all source locations. RE is the symmetric L1 norm difference; $r^2$ is Pearson $r^2$. Vectors are concatenated [LR; RC; VD] per source per sensor axis. Matched framework pairs (same bone model, BEM vs. FEM) are shaded.

| Model A | Model B | RE Med. | RE Max | $r^2$ Med. | $r^2$ Min |
|---|---|---|---|---|---|
| *Within-framework comparisons (BEM)* | | | | | |
| BEM Continuous | BEM Toroidal | 26.2% | 60.7% | 0.842 | 0.543 |
| BEM Continuous | BEM Realistic | 31.5% | 60.9% | 0.699 | 0.492 |
| BEM Toroidal | BEM Realistic | 9.0% | 24.6% | 0.978 | 0.817 |
| *Within-framework comparisons (FEM)* | | | | | |
| FEM Continuous | FEM Toroidal | 25.5% | 58.8% | 0.849 | 0.517 |
| FEM Continuous | FEM Realistic | 32.1% | 61.4% | 0.674 | 0.469 |
| FEM Toroidal | FEM Realistic | 11.5% | 26.5% | 0.966 | 0.774 |
| *Cross-framework matched pairs (BEM vs. FEM)* | | | | | |
| BEM Continuous | FEM Continuous | 2.5% | 7.3% | 0.998 | 0.975 |
| BEM Toroidal | FEM Toroidal | 3.1% | 6.2% | 0.998 | 0.990 |
| BEM Realistic | FEM Realistic | 1.9% | 5.6% | 0.999 | 0.992 |

#### Source-to-sensor distance analysis

Supplementary Figures S10 and S11 show maximum lead field amplitude as a function of closest source-to-sensor distance for the FEM and BEM realistic bone models respectively. In both frameworks, amplitude decreases monotonically with increasing source-to-sensor distance across all dipole orientations, with no systematic difference in the distance-amplitude relationship between BEM and FEM. Sources in the 230–300 mm region, where the localised BEM–FEM $r^2$ discrepancy for VD-oriented sources is observed, have source-to-sensor distances comparable to adjacent cord regions where BEM–FEM agreement is high. This confirms that the discrepancy cannot be attributed to proximity effects or amplitude magnitude differences between the two frameworks.

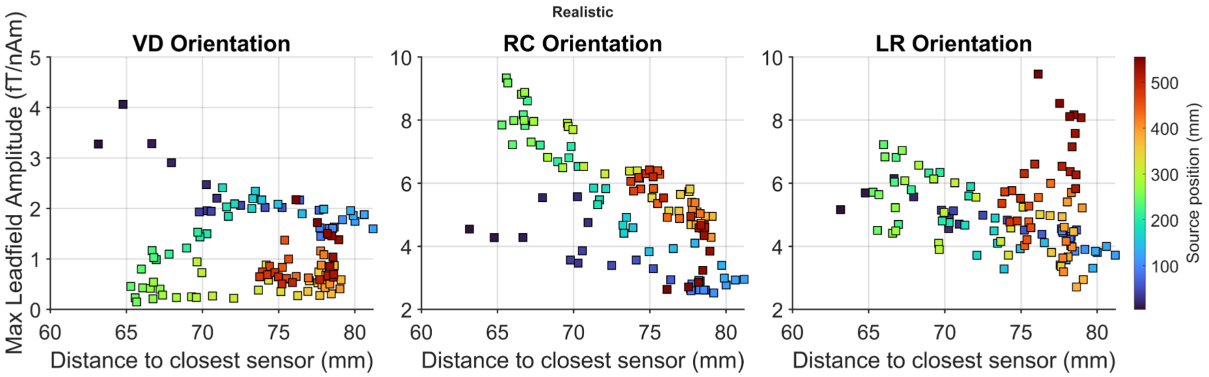


**Figure S10.** **Supplementary Figure S10 |** Maximum lead field amplitude (fT/nAm) as a function of closest source-to-sensor distance for FEM (MRI-derived realistic bone model), shown separately for each dipole orientation (VD: ventral–dorsal; RC: rostral–caudal; LR: left–right). Colour indicates source position along the spinal cord in millimetres, from the base of the brainstem (0 mm, dark blue) to the lumbar region (500 mm, dark red). The corresponding BEM result is provided in Supplementary Figure S11.

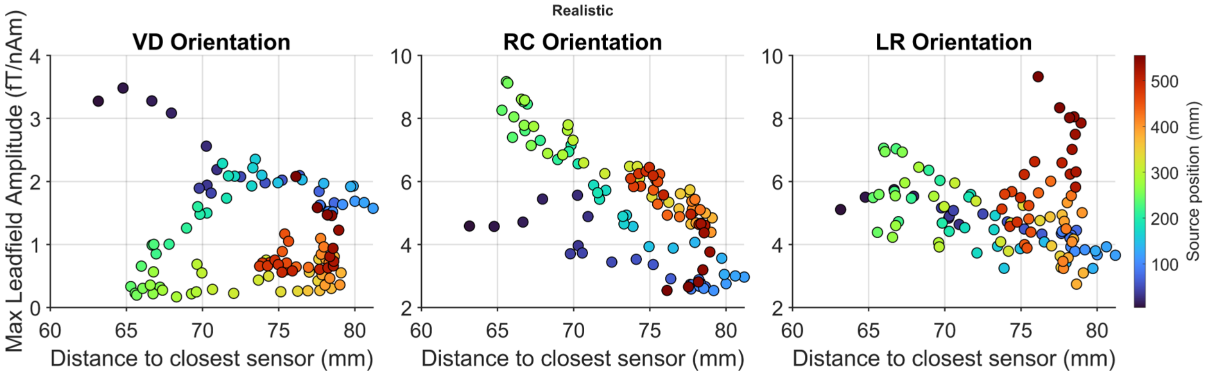


**Figure S11.** **Supplementary Figure S11 |** Maximum lead field amplitude (fT/nAm) as a function of closest source-to-sensor distance for BEM (MRI-derived realistic bone model). Format as in Supplementary Figure S10.

#### Organ removal: radial sensor axis

Supplementary Figure S12 shows source-wise $r^2$ and RE for the radial sensor axis (axis 3), comparing the BEM original against all organ-removal variants and the FEM original. For RC- and LR-oriented sources, all variants maintained high $r^2$ and low RE throughout the cord, consistent with the summary statistics in Supplementary Table S5. For VD-oriented sources, the localised $r^2$ reduction in the 230–300 mm region is present for all comparisons and is not resolved by organ removal. Removing both organs simultaneously worsened the BEM–FEM discrepancy relative to the individual-organ and original comparisons. Full per-orientation quantitative statistics for the 200–270 mm region are provided in Supplementary Table S6.

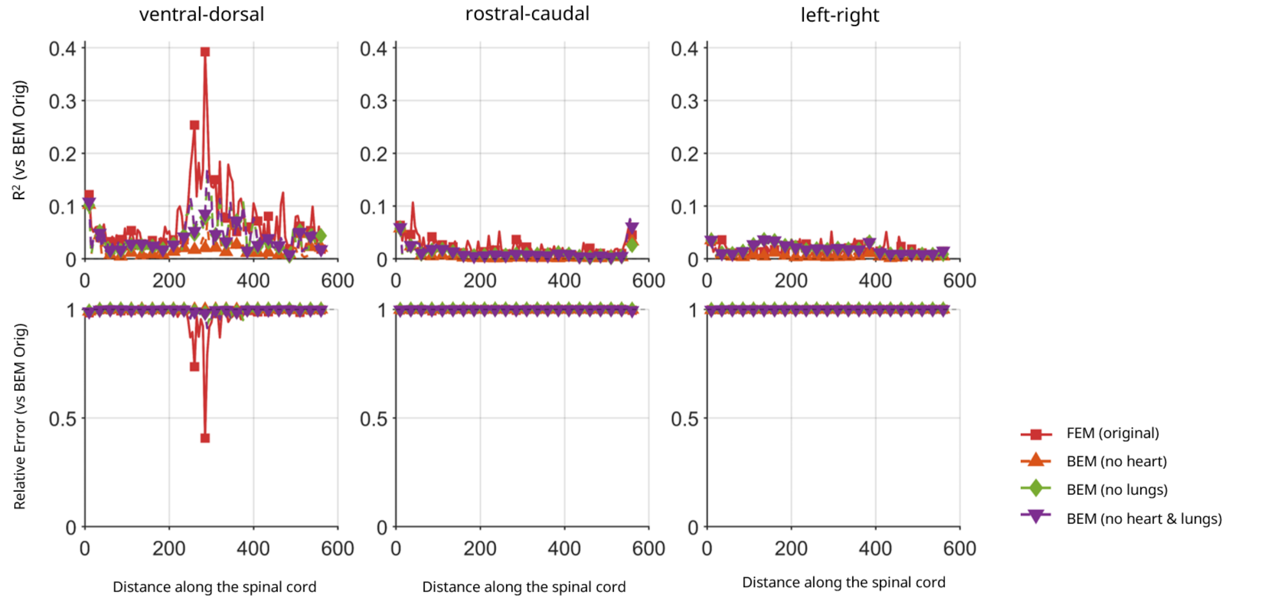


**Figure S12.** **Supplementary Figure S12** | Source-wise $r^2$ (top) and RE (bottom) for the radial sensor axis (axis 3), comparing the BEM original against BEM with the heart removed, BEM with the lungs removed, BEM with both removed, and FEM original. All comparisons are shown separately for each dipole orientation (VD: ventral–dorsal; RC: rostral–caudal; LR: left–right). Quantitative summary statistics are provided in Supplementary Tables S5 and S6.

#### Organ removal: tangential sensor axes

Supplementary Figures S13 and S14 show source-wise $r^2$ and RE for the two tangential sensor axes. The pattern on both tangential axes is consistent with the radial axis results in Supplementary Figure S12. For RC- and LR-oriented sources, all variants maintained $r^2$ above 0.986 and median RE below 6.5% across both tangential axes. For VD-oriented sources, the same localised reduction in $r^2$ is present in the 230–300 mm region, confirming that this effect is not specific to the radial field component but reflects the near-zero primary source contribution in this cord region and orientation across all measurable field components.

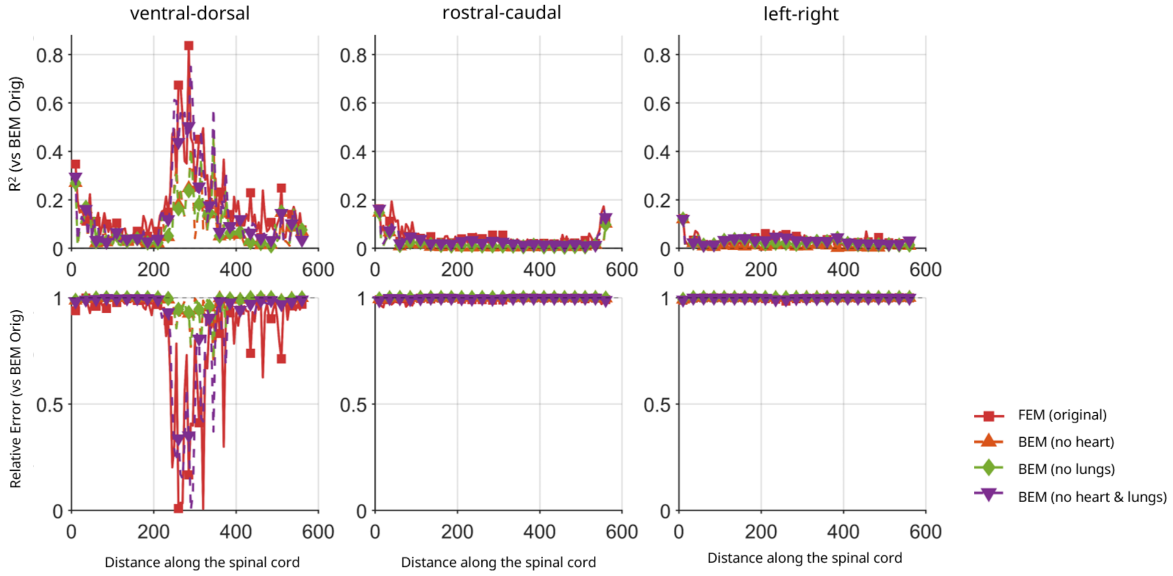


**Figure S13. Supplementary Figure S13 |** Source-wise $r^2$ (top) and RE (bottom) for sensor axis 1 (tangential), comparing the BEM original against BEM with the heart removed, BEM with the lungs removed, BEM with both removed, and FEM original. All comparisons are shown separately for each dipole orientation (VD: ventral–dorsal; RC: rostral–caudal; LR: left–right). Quantitative summary statistics are provided in Supplementary Table S5.

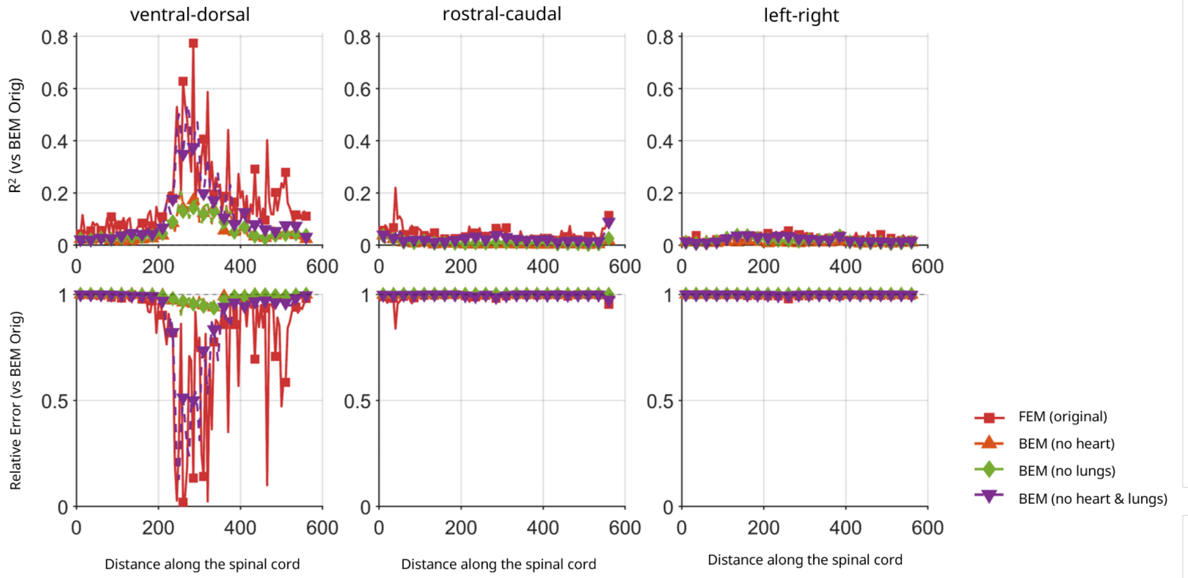


**Figure S14. Supplementary Figure S14 |** As Supplementary Figure S13 for sensor axis 2 (tangential).

#### Organ removal: cross-framework comparison per variant

Supplementary Figure S15 shows the BEM–FEM cross-framework comparison for each organ-removal variant on the radial sensor axis. For VD-oriented sources, removing the heart or lungs individually modestly reduced the BEM–FEM RE relative to the original (matched-pair median RE 10.6% and 19.5% respectively, versus 21.4% for the original; Supplementary Table S6), whilst removing both organs simultaneously increased it to a median RE of 27.1%, accompanied by a characteristic oscillation in $r^2$ between adjacent source positions ranging from 0.375 to values approaching unity. For RC-oriented sources, organ removal had minimal effect on BEM–FEM agreement, with all variants achieving median RE below 3.3% and $r^2$ above 0.997. For LR-oriented sources, median RE remained below 5.6% and $r^2$ above 0.993 across all variants, with the exception of the no-heart-and-lungs variant where a median amplitude difference of 26.3% was accompanied by $r^2 > 0.999$, indicating an amplitude scaling effect rather than topographic divergence.

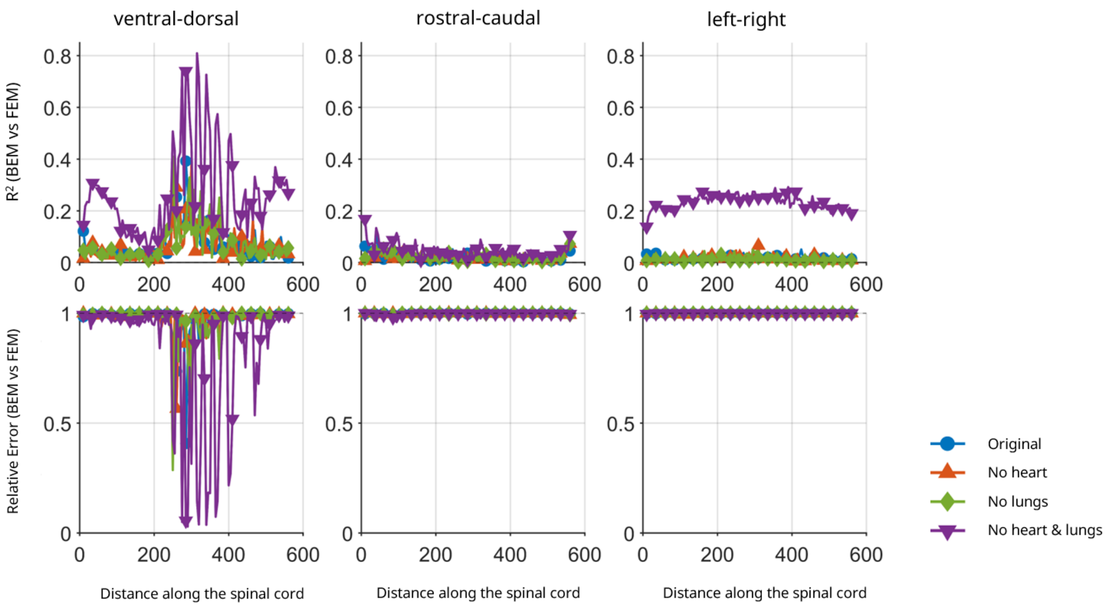


**Figure S15.** **Supplementary Figure S15 |** RE (top row) and $r^2$ (bottom row) comparing BEM versus FEM for each organ-removal variant (original, no heart, no lungs, no heart and lungs) on the radial sensor axis (axis 3). Each line represents the pairwise comparison between the BEM and FEM implementations of the same variant. Quantitative statistics for the 200–270 mm region are provided in Supplementary Table S6.

#### Organ removal: within-framework comparisons

Supplementary Figures S16 and S17 show source-wise RE and $r^2$ comparing organ-removal variants against their respective originals within each framework on the radial sensor axis. Within the BEM framework (Supplementary Figure S16), $r^2$ remained above 0.90 for all three dipole orientations throughout the cord. For RC- and LR-oriented sources, RE remained below 0.05 across all variants, with the no-heart-and-lungs variant producing the largest differences (median RE 2.89% for RC and 4.53% for LR). For VD-oriented sources, RE increased progressively between 200 and 400 mm, reaching a maximum of 0.07 for no heart, 0.10 for no lungs, and 0.15 for no heart and lungs, before returning toward baseline beyond 400 mm.

Within the FEM framework (Supplementary Figure S17), RC- and LR-oriented sources showed $r^2$ above 0.99 and RE below 0.10 for the no-heart and no-lungs variants. The no-heart-and-lungs variant produced substantially elevated amplitude differences for LR-oriented sources (median RE 26.4%), though $r^2$ remained above 0.998, indicating an amplitude scaling effect rather than topographic divergence. For VD-oriented sources, the FEM framework showed substantially greater sensitivity to organ removal than BEM, with the severity and persistence of the $r^2$ drop increasing with the extent of organ removal. The greater FEM sensitivity is consistent with the Discussion: FEM resolves internal volume current structure in greater detail and is therefore more sensitive to changes in the organ boundaries that govern secondary current pathways for near-radially oriented sources.

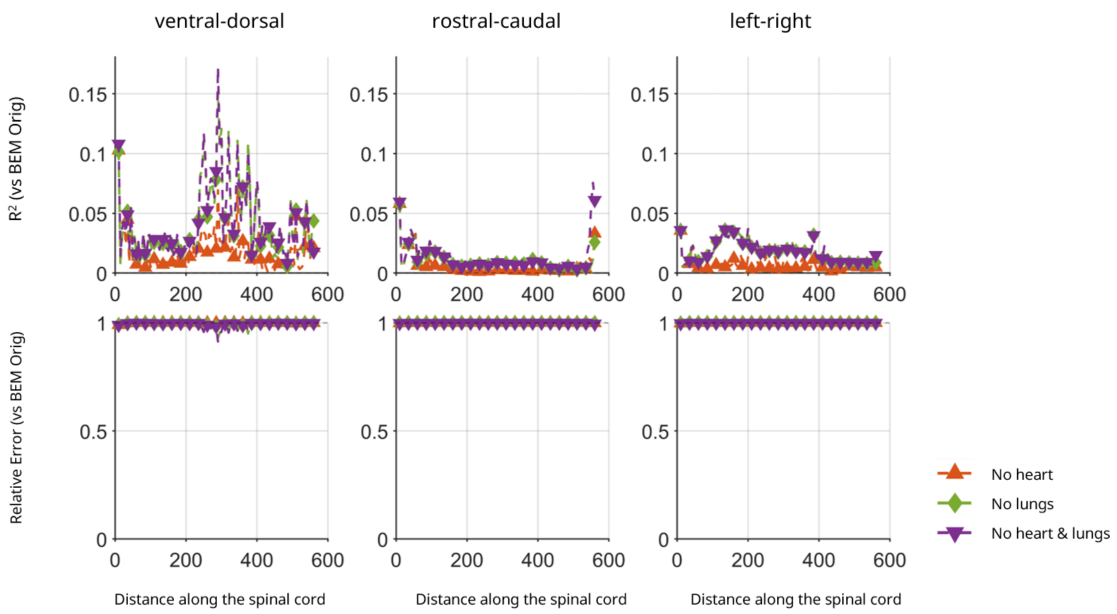


**Figure S16.** **Supplementary Figure S16 |** Source-wise RE (top) and $r^2$ (bottom) comparing each BEM organ-removal variant against the BEM original on the radial sensor axis (axis 3), shown separately for each dipole orientation (VD: ventral–dorsal; RC: rostral–caudal; LR: left–right). Quantitative statistics are provided in Supplementary Tables S5 and S6.

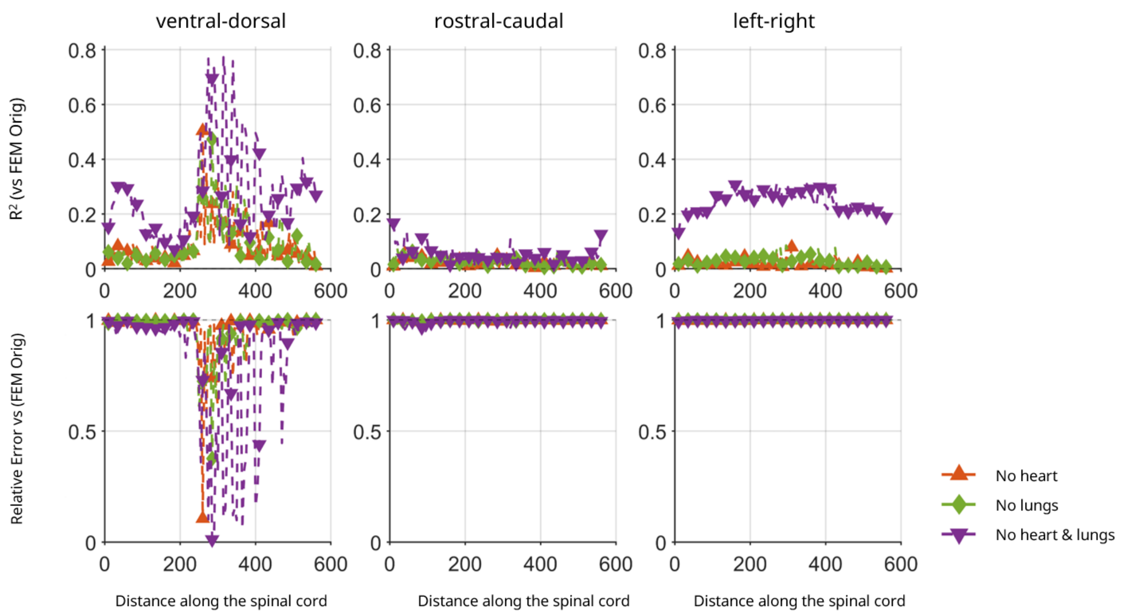


**Figure S17.** **Supplementary Figure S17 |** Source-wise RE (top) and $r^2$ (bottom) comparing each FEM organ-removal variant against the FEM original on the radial sensor axis (axis 3), shown separately for each dipole orientation (VD: ventral–dorsal; RC: rostral–caudal; LR: left–right). Quantitative statistics are provided in Supplementary Tables S5 and S6.

**Table S5. Supplementary Table S5 |** Summary RE and $r^2$ statistics comparing the BEM original against FEM and within-BEM organ-removal variants, across all three sensor axes. RE is computed as the symmetric L1 norm and $r^2$ as Pearson $r^2$, with lead field vectors concatenated across all three dipole orientations per source per sensor axis. Edge sources are excluded. Per-orientation statistics for the 200–270 mm region are provided in Supplementary Table S6.

| Sensor Axis | Comparison | RE Med (%) | RE Min (%) | RE Max (%) | $r^2$ Med | $r^2$ Min | $r^2$ Max |
|---|---|---|---|---|---|---|---|
| 4*Axis 1 | BEM Original vs. FEM Original | 3.65 | 1.54 | 17.64 | 0.9952 | 0.9700 | 0.9993 |
| | BEM Original vs. BEM No Heart | 1.20 | 0.51 | 15.49 | 0.9997 | 0.9907 | 1.0000 |
| | BEM Original vs. BEM No Lungs | 1.99 | 0.95 | 15.49 | 0.9993 | 0.9907 | 0.9999 |
| | BEM Original vs. BEM No Heart & Lungs | 3.01 | 1.68 | 16.54 | 0.9976 | 0.9860 | 0.9990 |
| 4*Axis 2 | BEM Original vs. FEM Original | 3.52 | 1.70 | 7.76 | 0.9956 | 0.9754 | 0.9992 |
| | BEM Original vs. BEM No Heart | 1.07 | 0.59 | 1.71 | 0.9997 | 0.9991 | 1.0000 |
| | BEM Original vs. BEM No Lungs | 1.71 | 0.67 | 3.10 | 0.9995 | 0.9978 | 1.0000 |
| | BEM Original vs. BEM No Heart & Lungs | 2.48 | 0.80 | 5.09 | 0.9983 | 0.9896 | 0.9999 |
| 4*Axis 3 (radial) | BEM Original vs. FEM Original | 1.93 | 0.68 | 5.61 | 0.9990 | 0.9918 | 0.9999 |
| | BEM Original vs. BEM No Heart | 0.52 | 0.17 | 5.29 | 0.9999 | 0.9970 | 1.0000 |
| | BEM Original vs. BEM No Lungs | 1.44 | 0.61 | 5.27 | 0.9996 | 0.9971 | 1.0000 |
| | BEM Original vs. BEM No Heart & Lungs | 1.48 | 0.70 | 5.45 | 0.9996 | 0.9969 | 0.9999 |

**Table S6. Supplementary Table S6 |** Pairwise RE and $r^2$ for the 200–270 mm cord region (sources 40–54), reported separately per dipole orientation for the radial sensor axis (axis 3). RE is the median symmetric L1 norm across sources in the region; $r^2$ Med and $r^2$ Min are the median and minimum squared Pearson correlation across sources. Entries marked * have their maximum amplitude difference at a source where absolute amplitude $<0.2$ fT/nAm and should be interpreted with caution. The no-heart-and-lungs BEM–FEM LR comparison (†) shows a large amplitude discrepancy (median RE 26.3%) driven by amplitude scaling rather than topographic divergence ($r^2 > 0.999$).

| Comparison | Ori. | RE Med (%) | $r^2$ Med | $r^2$ Min | Amp diff max (%) | Pos @ max (mm) |
|---|---|---|---|---|---|---|
| *BEM Original vs. FEM and organ-removal variants* | | | | | | |
| BEM Orig vs. FEM Orig | VD | 21.4 | 0.819 | 0.009 | 33.7 | 270 |
| BEM Orig vs. FEM Orig | RC | 3.2 | 0.994 | 0.989 | 6.5 | 260 |
| BEM Orig vs. FEM Orig | LR | 5.6 | 0.993 | 0.977 | 7.2 | 260 |
| BEM Orig vs. BEM No Heart | VD | 9.0 | 0.984 | 0.864 | 9.8 | 255 |
| BEM Orig vs. BEM No Heart | RC | 0.5 | 1.000 | 1.000 | 0.5 | 240 |
| BEM Orig vs. BEM No Heart | LR | 1.0 | 1.000 | 0.999 | 1.5 | 260 |
| BEM Orig vs. BEM No Lungs | VD | 9.0 | 0.986 | 0.856 | 8.4 | 255 |
| BEM Orig vs. BEM No Lungs | RC | 1.1 | 1.000 | 1.000 | 0.4 | 220 |
| BEM Orig vs. BEM No Lungs | LR | 2.1 | 1.000 | 0.998 | 3.7 | 200 |
| BEM Orig vs. BEM No H&L | VD | 15.5 | 0.906 | 0.150 | 59.8* | 255 |
| BEM Orig vs. BEM No H&L | RC | 2.9 | 0.997 | 0.994 | 3.1 | 270 |
| BEM Orig vs. BEM No H&L | LR | 4.5 | 0.993 | 0.984 | 5.7 | 200 |
| *BEM vs. FEM matched pairs (per variant)* | | | | | | |
| BEM Orig vs. FEM Orig | VD | 21.4 | 0.819 | 0.009 | 33.7 | 270 |
| BEM No Heart vs. FEM No Heart | VD | 10.6 | 0.940 | 0.113 | 43.4 | 270 |
| BEM No Lungs vs. FEM No Lungs | VD | 19.5 | 0.774 | 0.071 | 24.5 | 265 |
| BEM No H&L vs. FEM No H&L | VD | 27.1 | 0.875 | 0.375 | 65.5* | 255 |
| BEM Orig vs. FEM Orig | RC | 3.2 | 0.994 | 0.989 | 6.5 | 260 |
| BEM No Heart vs. FEM No Heart | RC | 3.1 | 0.996 | 0.986 | 8.0 | 240 |
| BEM No Lungs vs. FEM No Lungs | RC | 2.5 | 0.997 | 0.987 | 7.3 | 240 |
| BEM No H&L vs. FEM No H&L | RC | 3.3 | 0.998 | 0.997 | 5.2 | 250 |
| BEM Orig vs. FEM Orig | LR | 5.6 | 0.993 | 0.977 | 7.2 | 260 |
| BEM No Heart vs. FEM No Heart | LR | 4.0 | 0.993 | 0.985 | 6.2 | 245 |
| BEM No Lungs vs. FEM No Lungs | LR | 3.4 | 0.993 | 0.986 | 6.5 | 255 |

*Continued on next page*

*(Supplementary Table S6 continued)*

| Comparison | Ori. | RE Med (%) | $r^2$ Med | $r^2$ Min | Amp diff max (%) | Pos @ max (mm) |
|---|---|---|---|---|---|---|
| BEM No H&L vs. FEM No H&L | LR | 26.3† | 0.999 | 0.991 | 30.7 | 205 |
| *Within-BEM: organ-removal variants vs. BEM original* | | | | | | |
| BEM Orig vs. BEM No Heart | VD | 9.0 | 0.984 | 0.864 | 9.8 | 255 |
| BEM Orig vs. BEM No Lungs | VD | 9.0 | 0.986 | 0.856 | 8.4 | 255 |
| BEM Orig vs. BEM No H&L | VD | 15.5 | 0.906 | 0.150 | 59.8* | 255 |
| BEM Orig vs. BEM No Heart | RC | 0.5 | 1.000 | 1.000 | 0.5 | 240 |
| BEM Orig vs. BEM No Lungs | RC | 1.1 | 1.000 | 1.000 | 0.4 | 220 |
| BEM Orig vs. BEM No H&L | RC | 2.9 | 0.997 | 0.994 | 3.1 | 270 |
| BEM Orig vs. BEM No Heart | LR | 1.0 | 1.000 | 0.999 | 1.5 | 260 |
| BEM Orig vs. BEM No Lungs | LR | 2.1 | 1.000 | 0.998 | 3.7 | 200 |
| BEM Orig vs. BEM No H&L | LR | 4.5 | 0.993 | 0.984 | 5.7 | 200 |
| *Within-FEM: organ-removal variants vs. FEM original* | | | | | | |
| FEM Orig vs. FEM No Heart | VD | 14.2 | 0.960 | 0.029 | 52.3 | 265 |
| FEM Orig vs. FEM No Lungs | VD | 18.1 | 0.833 | 0.007 | 48.8 | 250 |
| FEM Orig vs. FEM No H&L | VD | 26.8 | 0.739 | 0.005 | 45.8 | 270 |
| FEM Orig vs. FEM No Heart | RC | 2.2 | 0.998 | 0.990 | 6.9 | 260 |
| FEM Orig vs. FEM No Lungs | RC | 3.0 | 0.997 | 0.992 | 6.2 | 210 |
| FEM Orig vs. FEM No H&L | RC | 4.1 | 0.997 | 0.990 | 6.9 | 260 |
| FEM Orig vs. FEM No Heart | LR | 2.5 | 0.998 | 0.990 | 9.8 | 260 |
| FEM Orig vs. FEM No Lungs | LR | 3.2 | 0.997 | 0.989 | 9.2 | 270 |
| FEM Orig vs. FEM No H&L | LR | 26.4† | 0.995 | 0.988 | 31.9 | 270 |
